# Inflammation perturbs hematopoiesis by remodeling specific compartments of the bone marrow niche

**DOI:** 10.1101/2024.09.12.612751

**Authors:** James W. Swann, Ruiyuan Zhang, Evgenia V. Verovskaya, Fernando J. Calero-Nieto, Xiaonan Wang, Melissa A. Proven, Peter T. Shyu, X. Edward Guo, Berthold Göttgens, Emmanuelle Passegué

## Abstract

Hematopoietic stem and progenitor cells (HSPC) are regulated by interactions with stromal cells in the bone marrow (BM) cavity, which can be segregated into two spatially defined central marrow (CM) and endosteal (Endo) compartments. However, the importance of this spatial compartmentalization for BM responses to inflammation and neoplasia remains largely unknown. Here, we extensively validate a combination of scRNA-seq profiling and matching flow cytometry isolation that reproducibly identifies 7 key CM and Endo populations across mouse strains and accurately surveys both niche locations. We demonstrate that different perturbations exert specific effects on different compartments, with type I interferon responses causing CM mesenchymal stromal cells to adopt an inflammatory phenotype associated with overproduction of chemokines modulating local monocyte dynamics in the surrounding microenvironment. Our results provide a comprehensive method for molecular and functional stromal characterization and highlight the importance of altered stomal cell activity in regulating hematopoietic responses to inflammatory challenges.

## Introduction

The hematopoietic system maintains a stable output of mature blood cells under homeostatic conditions, but hematopoietic stem and progenitor cells (HSPC) also need to respond to demands by increasing production of specific blood lineages (Swann et al., 2024). Accumulating evidence shows that overall hematopoietic responses are frequently attributable to changes in number and function of both HSPCs and stromal cells located together in the marrow cavity, highlighting the importance of stromal cells in modulating HSPC function upon a range of challenges, including bacterial and viral infections (Boettcher et al., 2014; Isringhausen et al., 2021), aging (Mitchell et al., 2023), and leukemia (Schepers et al., 2013; 2015). However, despite growing recognition that interacting hematopoietic and stromal cells form the functional unit in marrow responses, the manner in which distinct spatial compartments of the bone marrow (BM) niche respond to different stimuli has received little attention to date. Gaining insights into these regionalized ecosystems will be instrumental in understanding how the function of specific stromal cell populations can be modulated to mitigate inflammatory responses and disease development.

Adapting blood cell production to demand relies on interactions between HSPCs and stromal cells in the BM niche, including endothelial cells (EC), mesenchymal stromal cells (MSC), and a range of mesenchymal progenitors (MPr) including osteoblast progenitors (OPr), chrondroblast progenitors (CPr), and fibroblast progenitors (FPr) (Pinho & Frenette, 2019; Woods & Guezguez, 2021). Stromal cells regulate HSPCs through direct cell-cell contact and by producing soluble ligands that diffuse in the surrounding BM microenvironment. For instance, stromal-derived interleukin (IL)-7 is essential for common lymphoid progenitor (CLP) development (Zhu et al., 2007; Wu et al., 2008), whereas production of trophic factors like CXCL12 and stem cell factor (SCF or Kit ligand) by leptin receptor (LepR)-expressing MSCs (MSC-L) and BM ECs is required for retention of hematopoietic stem cells (HSC) in the BM niche, as well as for enforcing HSC quiescence to preserve lifelong self-renewal (Ding et al., 2012; Ding & Morrison, 2013; Greenbaum et al., 2013; Asada et al., 2017). Direct contacts between HSPCs and stromal cells mediated by integrins like the very late antigen (VLA)-4 and VLA-5 are also essential for long-term survival of hematopoietic cells *ex vivo* (Jung et al., 2005), and similar interactions between HSPCs, stromal cells, and extracellular matrix proteins (ECM) organize the BM niche into distinct spatial domains with specialized functions (Baccin et al., 2020).

BM stromal cells have been characterized by imaging of bone sections, flow cytometry of dissociated tissue, and more recently using emerging single cell RNA sequencing (scRNA-seq) and spatial transcriptomic technologies. These complementary approaches describe a BM niche composed of two principal spatial compartments: the soft central marrow (CM) niche, which is vascularized by an extensive network of vacuolized sinusoids, and the collagen rich endosteal (Endo) niche that lines the inner bone surfaces and is supplied with blood by arterioles (Kusumbe et al., 2014; Itkin et al., 2016; Ramalingam et al., 2021). Different HSPC populations are selectively enriched in either compartment, with CLPs and a subset of deeply quiescent HSCs being preferentially located at the endosteum (Lo Celso et al., 2009; Ding & Morrison, 2013), and most HSCs, multipotent progenitors (MPP), granulocyte macrophage progenitors (GMP), and megakaryocytes (Mk) being distributed in the central marrow, sometimes organized along sinusoidal vessels in maturing production lines (Zhang et al., 2021). Despite this clear segregation in cell distribution and functional hematopoietic activity, most investigatory methods do not permit separate profiling of stromal cells originating from these two compartments, hampering investigations of diseases like cancer or inflammation that simultaneously affect both stromal and hematopoietic compartments. While this limitation can be addressed to some extent using global approaches that sample all cells, such as scRNA-seq, there is a need for a widely applicable, cost-effective technique that permits reliable, prospective isolation of functionally validated stromal populations from both Endo and CM locations to facilitate investigation of BM niche perturbations across mouse strains.

Here, we describe a refined methodology to separately isolate and profile stromal cells of CM and Endo spatial niches using complementary scRNA-seq profiling and flow cytometry analyses. We validate this method extensively against orthogonal studies of the BM niche and establish its broad applicability in profiling stromal populations in the face of diverse challenges to understand how changes in specific spatial compartments shape the response of the hematopoietic system. In particular, we identify MSC-L as a critical first responder to type I interferons, licensing this population to regulate myeloid responses in the CM compartment of the BM niche.

## Results

### Profiling BM spatial compartments at single cell resolution

We developed a method to separately isolate niche compartments by enzymatic digestion of either a single intact marrow plug flushed from a mouse femur to obtain the CM preparation, or bone chips collected from eight bones (2 femurs, 2 tibiae, 2 humeri, and 2 hemipelves) crushed together and washed of non-adherent BM cells to obtain the Endo preparation (**Fig. 1a**). Profiling of HSPCs remaining at the endosteum after flushing the marrow plug and in the marrow plug itself confirmed the importance of spatial separation, with the previously reported preferential enrichment of HSCs and Mk progenitors (MkP) in the CM preparation, and CLPs in the Endo preparation (Ding & Morrison, 2013; Zhang et al., 2021) (**Extended Data Fig. 1a, Supplementary Fig. 1**). To identify which stromal cells were present at each location, we performed droplet-based 10X Genomics scRNA-seq on flow-isolated Ter119^-^/CD45^-^ CM and Endo cells from young wild type (WT) C57Bl/6 mice in three independent experiments to establish a 10X stroma map (**Supplementary Fig. 2a-c**). Analysis of the integrated dataset revealed little overlap of cells originating from either location in Louvain clusters, with only a few mesenchymal and endothelial clusters showing mixed Endo/CM origin (**Fig. 1b; Supplementary Fig. 2b**). However, significant and highly variable contamination by hematopoietic cells was observed across experiments, despite similar initial flow cytometry sorting gates (**Fig. 1c; Extended Data Fig. 1b; Supplementary Fig. 2d; Extended Data Table 1**). Both CM and Endo fractions contained CD45^+^ leukocytes in variable numbers that were primarily granulocytes, as well as CD45^-^ erythroid progenitors (EryP) and occasionally MkPs and other HSPCs. EryPs have already been described as typical contaminants of stromal preparations (Boulais et al., 2018), and differences in the degree of hematopoietic cell contamination is likely to contribute to the variability in cell type recovery observed among different investigators (Baryawno et al., 2019; Baccin et al., 2020; Mitchell et al., 2023) (**Extended Data Fig. 1c)**. We also confirmed the importance of lysing red blood cells (RBC) in stromal preparations to limit the number of contaminating EryP (**Extended Data Fig. 1c)**, which can be further complemented by selectively depleting cells expressing the erythroid marker CD71 prior to scRNA-seq analyses (Baryawno et al., 2019). Despite variations in contaminating hematopoietic cells, recovery of stromal cells was consistent in our three separate datasets, with all major cell clusters represented across replicates (**Supplementary Fig. 2b**). This establishes the repeatable nature of our method to isolate stromal cells from different spatial locations in the BM niche.

**Figure 1.**
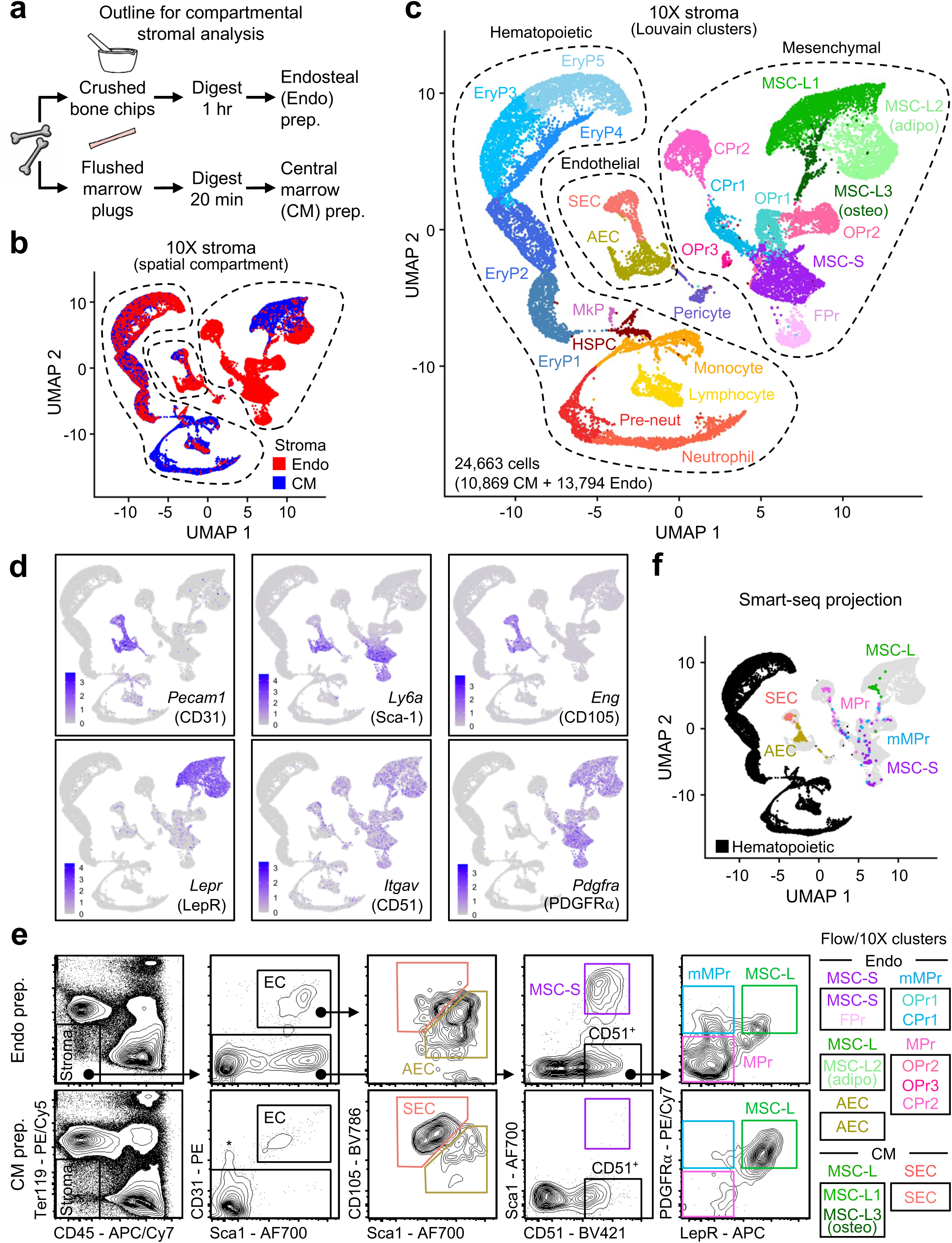
Stromal cells are arranged in two distinct spatial compartments. **a,** Isolation strategy for stromal cells in the central marrow (CM) and endosteal (Endo) fractions. **B,** Uniform manifold and approximation projection (UMAP) of integrated 10X single cell RNA sequencing (scRNA-seq) stroma datasets derived from n = 3 independent experiments profiling CM (blue) and Endo (red) fractions. **C,** UMAP showing Louvain clusters with number of analyzed cells, separation of mesenchymal and endothelial cells from hematopoietic contaminants, and cluster nomenclature. AEC: arterial endothelial cells, SEC: sinusoidal endothelial cell, MSC-S: Sca-1^+^ mesenchymal stromal cell (MSC), OPr: osteoblast progenitor, CPr: chondroblast progenitor, FPr: fibroblast progenitor, MSC-L: leptin receptor (LepR)^+^ MSC (adipo: adipodegnic, osteo: osteogenic), EryP: erythroid progenitor, MkP: megakaryocyte progenitor, HSPC: hematopoietic stem and progenitor cells, Pre-neut, pre-neutrophil. **D,** Feature plots showing expression of indicated genes in the 10X stroma dataset. **E,** Representative flow cytometry plots showing gating of key stromal populations in Endo and CM preparations (prep.) with flow/10X cluster correspondence. (m)MPr: (multipotent) mesenchymal progenitor. * denote contamination by EryP expressing low levels of CD31 marker. **F,** Projection of Smart-seq stromal cell transcriptomes onto the 10X stroma dataset.

### Spatial compartmentalization of the BM niche can be resolved by flow cytometry

We next investigated the identity of distinct Louvain clusters identified in our combined Endo/CM 10X stroma map (**Fig. 1c**). Among the EC clusters, we annotated a group of arteriolar endothelial cells (AEC) of almost exclusively Endo origin (96%) and a group of sinusoidal endothelial cells (SEC) of mixed origin (59% CM, 41% Endo) (**Extended Data Table 1**). We also confirmed the presence of BM pericytes (Baryawno et al., 2019) that were predominantly of Endo origin (93%). Gene expression analyses showed broad expression of *Pecam1* (CD31), with opposite expression patterns for *Ly6a* (Sca-1) and *Eng* (CD105) in AECs vs. SECs (**Fig. 1d, Extended Data Fig. 1d**). This was consistent with previously described flow cytometry analyses of these two endothelial populations in the Ter119^-^/CD45^-^ stromal fraction (Helbling et al., 2019; Mitchell et al., 2023), with AECs identified as CD31^+^/Sca-1^bright^/CD105^low^ cells and found preferentially in the Endo preparation (85.9 ±1.6% of total ECs), and SECs as CD31^+^/Sca-1^low^/CD105^bright^ cells enriched mainly in the CM preparation (70.7 ±7.2% of total ECs) (**Fig. 1e; Extended Data Fig. 1e**). We also performed 10X scRNA-seq on AECs and SECs isolated from both CM and Endo preparations and projected these flow-purified populations onto our 10X stroma map based on nearest neighbor embedding to confirm their identification (**Extended Data Fig. 2a**). Interestingly, while Endo SECs were indistinguishable from the more abundant CM SECs, the small numbers of CM AECs were rather distinct from Endo AECs and displayed higher *Emcn* expression (**Extended Data Fig. 2b**), likely representing the transitional Endomucin high (Emcn^hi^) type-H vessels described by imaging (Kusumbe et al., 2014). This demonstrated that the two main EC groups found in the marrow cavity can be captured effectively by flow cytometry and investigated for molecular heterogeneity by scRNA-seq analyses.

Among the mesenchymal clusters, we first annotated 3 clusters of MSC-Ls that were of mixed origin, with MSC-L1 and MSC-L3 composed predominantly of CM cells (88% and 67%, respectively) and MSC-L2 mostly formed of Endo cells (88%) (**Fig. 1b,c; Extended Data Table 1**). Gene expression analyses showed expression of *Cxcl12* (CXCL12), *Itgav* (CD51), and *Pdgfra* (PDGFR-α) across all 3 clusters, but suggested a more committed adipogenic fate in MSC-L2 as indicated by higher *Apoe* and *Adipoq* expression, and a strong bias towards osteogenic fate in MSC-L3 as shown by *Alpl* and *Bglap* expression and by their transcriptomic similarity to Endo-derived OPr2 (**Fig. 1c; Extended Data Fig. 2c**). This was also consistent with our flow cytometry isolation of MSC-Ls as Sca-1^-^/CD51^+^/PDGFRα^+^/LepR^+^ cells in the non-endothelial CD31^-^/Ter119^-^/CD45^-^ fraction at both locations, with Endo MSC-Ls representing a small subset of Endo mesenchymal cells (2.4 ± 1.0% of CD51^+^ cells) and CM MSC-Ls forming the vast majority of CM mesenchymal cells (88.1 ± 4.3% of CD51^+^ cells) (**Fig. 1e; Extended Data Fig. 1e**). This distribution was also validated by projection of flow-purified MSC-Ls isolated from both CM and Endo preparations and analyzed by 10X scRNA-seq onto our 10X stroma map, which confirmed that MSC-L2 was primarily composed of Endo MSC-Ls, whereas uncommitted MSC-L1 was exclusively derived from CM MSC-Ls (**Extended Data Fig. 2a**). We also annotated a large mesenchymal group composed of 7 *Itgav*-expressing clusters that was of exclusive Endo origin. Co-expression of *Ly6a* and *Pdgfra* identified Sca-1-expressing MSCs (MSC-S), which closely resemble previously defined PαS cells (Helbling et al., 2019; Pisterzi et al., 2023), and a group of fibroblast progenitors (FPr) identified based on expression of markers like *Sema3c* (**Extended Data Fig. 2d; Extended Data Table 1**). Further populations were separated into *Pdgfra^+^*multipotent mesenchymal progenitors (mMPr) and more committed *Pdgfra^-^*mesenchymal progenitors (MPr) with either a chondroblastic (CPr1/2) or osteoblastic (OPr1/2/3) bias (**Fig. 1d; Extended Data Fig. 2d; Extended Data Table 1**). This again was consistent with our flow cytometry isolation of MSC-S (including FPr) as Sca-1^+^/CD51^+^/PDGFRα^+^/LepR^-^ cells, mMPr (representing OPr1/CPr1) as Sca-1^-^/CD51^+^/PDGFRα^+^/LepR^lo^ cells, and MPr (covering OPr2/OPr3/CPr2) as Sca-1^-^/CD51^+^/PDGFRα^−^/LepR^-^ cells in the non-endothelial CD31^-^/Ter119^-^/CD45^-^ fraction of the Endo preparation (13.6 ± 8.6% of endosteal CD51^+^ cells for MSC-S, 4.1 ± 1.8% for mMPr, and 62.3 ± 12.8% for MPr) (**Fig. 1e; Extended Data Fig. 1e; Supplementary Fig. 2e,f**). In contrast, the non-endothelial CD31^-^/Ter119^-^/CD45^-^ fraction of the CM preparation was completely devoid of MSC-S and only had small numbers of mesenchymal progenitors (5.0 ± 2.7% of CM CD51^+^ cells for mMPr, and 5.7 ± 3.3% for MPr). Finally, we found that isolation of mesenchymal cells critically depends on enzymatic tissue digestion, and that using Collagenase I by itself provided excellent release and recovery of tissue-resident cells at both locations compared to other described extraction methods (**Extended Data Fig. 2e**). Taken together, these results indicate that the entire mesenchymal compartment can be profiled by flow cytometry, with added molecular resolution of population heterogeneity provided by scRNA-seq analyses. They also indicate that most of Endo populations (AEC, MSC-S, mMPr, MPr) remain associated with the bone chips during crushing, while the majority of CM populations (SEC, MSC-L) are flushed in the marrow plug, with cross compartment representation seen only in ECs and different adipogenic and osteogenic subsets of MSC-Ls.

To further validate the broad applicability of our flow cytometry scheme, we used our previously published plate-based Smart-seq scRNA-seq analyses of the major Endo and CM populations to project onto our 10X stroma map (**Fig. 1f; Extended Data Fig. 3a,b; Extended Data Table 2**) (Mitchell et al., 2023). We confirmed representation of all major stromal cell types in our dataset, with pseudobulk gene expression profiles for each cluster correlating strongly with the related populations defined by flow cytometry (**Extended Data Fig. 3c**). We also projected our Smart-seq transcriptomes onto three independent 10X datasets that used different approaches for stromal cell isolation (Baryawno et al., 2019; Tikhonova et al., 2019; Baccin et al., 2020), finding in each case that all major stromal populations were identified, albeit with considerable variation in naming conventions (**Extended Data Fig. 4a,b**). Collectively, these analyses demonstrate that all major stromal cell types can be identified by a simple flow cytometry method, permitting quantification and prospective isolation of cell populations that are spatially segregated into the CM and Endo compartments of the murine BM niche.

### Endothelial cells form two vascular niches with distinct functional properties

To determine whether flow-isolated CM SECs and Endo AECs correspond to similar sinusoidal and arterial cell types described in previous molecular and imaging studies, we first compared gene expression from our Smart-seq profiling (**Extended Data Table 2**) (Mitchell et al., 2023). Numerous differentially regulated genes (DEG) were identified, including several key markers used previously to distinguish arterial and sinusoidal ECs (Xu et al., 2018; Tikhonova et al., 2019; Baryawno et al., 2019) (**Fig. 2a,b**). Geneset enrichment analysis (GSEA) of genes upregulated in SECs using GO biological pathways revealed enrichment for VEGF signaling (**Fig. 2c**), which was directly confirmed by flow cytometry analyses of *Vegfr3-Yfp* reporter mice (Calvo et al., 2011), with higher expression of VEGFR3 in SECs compared to AECs (**Extended Data Fig. 5a**). We also found significant enrichment for pathways associated with leukocyte interaction in SECs, with higher expression of selectins and integrins (**Fig. 2d**), consistent with sinusoids being key interchange sites for cells moving between the BM and blood (Frenette et al., 1998). Accordingly, and in agreement with a previous report (Ramalingam et al., 2017), we showed that Endomucin, which regulates endothelial:leukocyte interactions and VEGF signaling (Zhang et al., 2020), was expressed by all SECs and to a much higher extent than AECs at the protein level (**Extended Data Fig. 5b**). Finally, SECs were enriched for pathways related to vascular permeability and were significantly more permeable to fluorescent dragon green beads (DGB) injected intravenously (**Fig. 2e,f**). Conversely, AECs were enriched for DEG signatures related to endothelial integrity (**Fig. 2g**), with a subpopulation of AECs selectively expressing the gap junction protein connexin 40 (CX40) encoded by *Gja5* which we directly confirmed by flow cytometry using *Cx40-Gfp* reporter mice (Miquerol et al., 2004) (**Extended Data Fig. 5c**). AECs were also enriched for Notch signaling pathway genes, displaying greater expression of Notch receptors and downstream response elements such as *Hey1* and *Hes1* compared to SECs, which we directly confirmed by flow cytometry using *Hes1-Gfp* reporter mice (Fre et al., 2011) (**Fig. 2h,i**). This enrichment for Notch signaling was confined to AECs and supported the role of the Notch ligand Dll4 in controlling CLP numbers in the same endosteal BM compartment (Tikhonova et al., 2019). In addition, we confirmed that ∼40% of AECs express high Nestin (Nes) level in *Nestin-Gfp* reporter mice (Mignone et al., 2004) (**Extended Data Fig. 5d**), although the transcriptomes of Nes^hi^ and Nes^lo^ AECs both correlated strongly with our Smart-seq profiled AECs (**Extended Data Fig. 5e**), consistent with previous data showing few transcriptional differences between AECs with different levels of Nestin expression (Xu et al., 2018). Collectively, these investigations provide extensive validation that flow-isolated SECs and AECs correspond phenotypically and functionally with similar arterial and sinusoidal EC populations described in previous studies. However, Sca-1 expression, a key identifying marker of AECs, is notoriously labile during inflammation and may be negligibly expressed in mice with the Ly6a1 haplotype (Spangrude & Brooks, 1993). We therefore searched for other cell surface markers that might distinguish AECs from SECs, finding that CD34 was expressed at significantly higher levels in AECs (**Extended Data Fig. 5f**). Accordingly, we found that CD34 effectively distinguished AECs and SECs in BALB/C mice, which possess the Ly6a1 haplotype and do not express Sca-1 on AECs (**Extended Data Fig. 5g**). The addition of CD34 therefore extends the applicability of our flow cytometry scheme to identify AECs and SECs across mouse strains.

**Figure 2.**
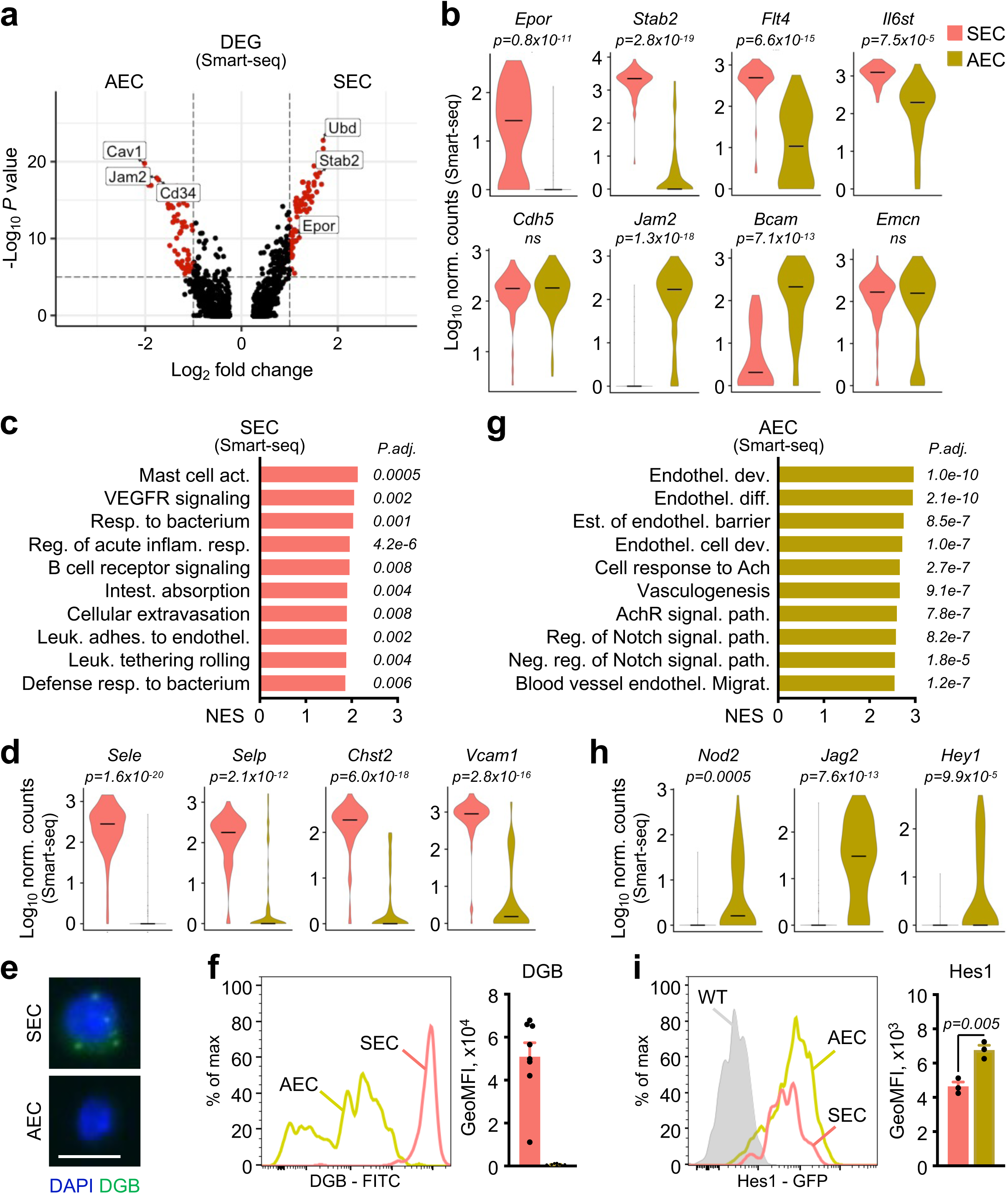
BM niche contains two functionally distinct types of endothelial cell. **A,** Volcano plot showing significantly differentially expressed genes between arterial (AEC) and sinusoidal (SEC) endothelial cells profiled by plate-based Smart-seq scRNA-seq. Red points show genes with log_2_ fold change >|1| and adjusted p value <10^-5^. **B,** Expression of signature genes associated with AEC and SEC identity. **C-f**, SEC characterization: (c) geneset enrichment analysis (GSEA) of selectively enriched GO biological pathways (NES: normalized enrichment score, P.adj.: adjusted *P values*); (d) examples of preferentially expressed genes; (e) representative images of dragon green bead (DEG) uptake (scale bar: 5 μm); and (f) representative flow cytometry plots (left) and quantification of geometric mean fluorescent intensity (GeoMFI, right) of DGB uptake. **g-i,** AEC characterization: (g) GSEA of enriched GO biological pathways; (h) examples of preferentially expressed genes; and (i) representative flow cytometry plots (left) and quantification of GeoMFI (right) of *Hes1*-GFP expression. Data in (b), (d), and (h) are violin plots of Log10 normalized (norm.) Smart-seq counts; *P. values*, Wilcoxon rank sum test. Data in (f) and (i) are means ± S.D. with points showing values for individual mice; *P. values*, Student’s t test. For (c) and (g), *P. values* are derived from permutation tests.

### Mesenchymal lineages are developmentally and spatially segregated in the BM niche

To understand the functional roles of flow-isolated mesenchymal populations, we isolated each major Endo population (MSC-S, mMPr, and MPr cells) as well as CM MSC-Ls for *ex vivo* differentiation assays. Although all mesenchymal populations could generate fibroblast colony forming units (CFU-F) (**Fig. 3a**), these were observed much more frequently from Endo populations according to their differentiation gradient (MSC-S > mMPr > MPr) than from CM MSC-Ls, which produce CFU-F infrequently and only under hypoxic conditions (5% O_2_) and in the presence of a ROCK kinase inhibitor (iROCK) to modulate sensing of the plastic substrate, as previously reported (Khatiwala et al., 2006; Yue et al., 2016) (**Fig. 3b**). Interestingly, in these conditions and upon osteoblastic differentiation, MSC-Ls formed the largest alkaline phosphatase (ALP) or von Kossa positive colonies compared to all Endo populations, which in terms of frequency were equivalent to MSC-Ls but formed smaller colonies (**Fig. 3b,d**). This confirms the potent bone-forming potential of MSC-Ls, which maintain bone in adult mice and are essential for repairing bone fractures (Zhou et al., 2014), and further illustrates the osteogenic potential of MSC-Ss and their (m)MPr derivatives that was initially demonstrated by transplantation (Morikawa et al., 2009). Lastly, only MSC-S and MSC-L cells had the capacity to form colonies under conditions promoting adipogenic differentiation (**Fig. 3e**), suggesting that this lineage potential is lost in more committed (m)MPr derivatives, consistent with the established role of MSC-Ls in generating most adipocytes *in vivo* at steady state and during regeneration (Zhou et al., 2014; Hirakawa et al., 2023). Altogether, these functional results confirm the multipotent nature of MSC-Ss and MSC-Ls, yet with non-overlapping biases towards distinct lineages, and highlight the more committed fates of (m)MPr cells in producing only endosteal lineage cells.

**Figure 3.**
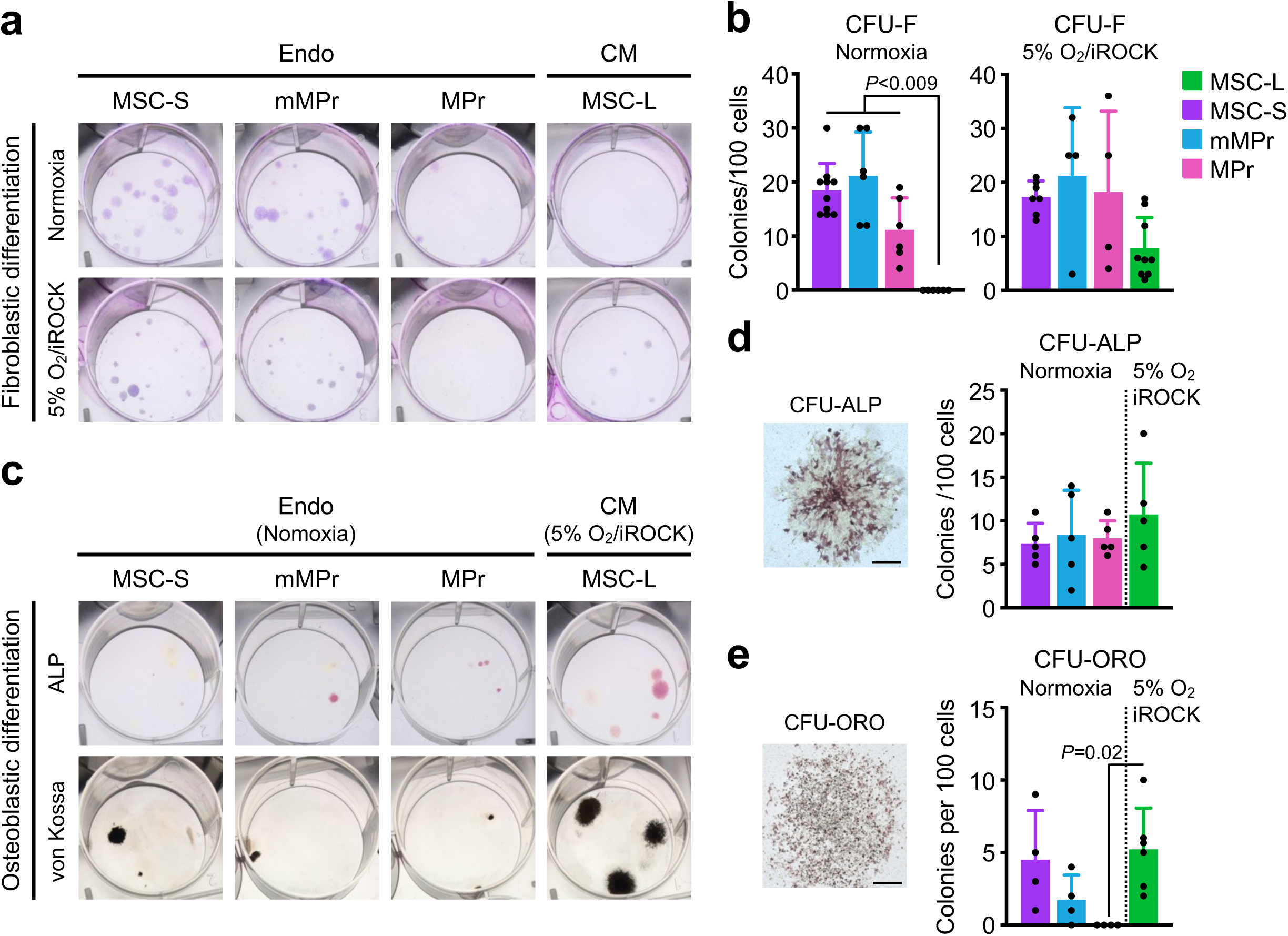
Bone forming-potential is present in CM and Endo compartments. **a-b,** Representative images (a) and quantification (b) of colony forming unit-fibroblast (CFU-F) obtained from the indicated Endo and CM populations. FACS-sorted cells (200 cells/35-mm well) were cultured for 11 days in normoxia or for 8 days in 5% O_2_ hypoxia with a ROCK kinase inhibitor (irOCK). **c-d,** Representative images (c) and quantification (d) of alkaline phosphatase expressing colonies (CFU-ALP) and calcifying colonies (von Kossa) obtained after 9 days (CFU-ALP) and 21 days (von Kossa) of induced osteoblastic differentiation of CFU-F colonies. A representative photomicrograph of a CFU-ALP colony (scale bar: 1 mm) is shown on the left in (d). **e,** Quantification of oil red-O expressing colony (CFU-ORO) obtained after 8 days (5% O_2_/iROCK) and 11 days (normoxia) of induced adipogenic differentiation of CFU-F colonies, with a representative photomicrograph of a CFU-ORO colony (scale bar: 1 mm) shown on the left. Data are means ± S.D. with points showing values for individual mice; *P. values*, one-way ANOVA with Tukey *post-hoc* test.

Although informative, *ex vivo* assays only define the differentiation potential of cell populations rather than the likelihood of their lineage commitment *in situ*, which is further complicated by the known plasticity of mesenchymal lineage cells (Herzog et al., 2003). We therefore sought to cross-reference our findings with molecular data capturing developmental profiles of mesenchymal cells in the BM niche that we obtained by integrating data from three independent published 10X scRNA-seq profiling studies (Baryawno et al., 2019; Baccin et al., 2020; Zhong et al., 2020) to create a mesenchymal atlas in which we could identify all major mesenchymal cell states (**Fig. 4a, Extended Data Fig. 6a-d**). Importantly, integrating datasets with differences in cell preparation maximized our coverage of cell types, allowing us to assess developmental relationships with greater resolution than in previous studies (Wolock et al., 2019) (**Extended Data Fig. 6e**). We then used published gene signatures to annotate two sources of BM mesenchymal cells: Sca-1^+^ MSCs and LepR^+^ MSCs (Zhong et al., 2020; Pisterzi et al., 2023) (**Fig. 4b; Extended Data Table 3**). Using these populations as starting nodes for pseudotime analysis (Trapnell et al., 2014), we inferred that Sca-1^+^ MSCs generated fibroblasts, chondroblasts, and osteoblasts, whereas LepR^+^ MSCs were upstream of osteoblasts and adipocytes (**Fig. 4a,b**). Importantly, projection of our 10X stroma map onto this 10X mesenchymal atlas confirmed that LepR^+^ MSCs and their progeny were present in both CM and Endo compartments, whereas Sca-1^+^ MSCs and their downstream lineages were confined to the Endo compartment (**Extended Data Fig. 6f**). We also showed strong overlap between the transcriptomes of Sca-1^+^ MSCs and LepR^+^ MSCs with our phenotypically defined and flow-isolated MSC-S and MSC-L populations, respectively (**Fig. 4c**). In agreement with their proposed potentiality, we found significant enrichment for GO pathways related to bone formation and mesenchymal development in MSC-Ls, and to fibroblast differentiation and extracellular matrix production in MSC-Ss (**Fig. 4d**). Our annotation of these two distinct sources of BM mesenchymal cells largely aligns with previous studies employing different methodologies. Thus, although both MSC-S and MSC-L cells expressed *Prx1* in *Prx1-tdTomato* reporter mice consistent with their mesenchymal origins (Logan et al., 2002) (**Extended Data Fig. 7**), only MSC-Ls overlapped with Nes^+^ cells by flow cytometry (Greenbaum et al., 2013; Kunisaki et al., 2013; Zhou et al., 2014) (**Extended Data Fig. 7**) and strongly correlated with the transcriptomes of cells selected based on expression of CXCL12 and Kit ligand (**Extended Data Fig. 8a**). Conversely, MSC-Ss, which were still identifiable using Sca-1 in mice with the Ly6a1 haplotype (**Extended Data Fig. 8b**), had a similar transcriptome to PαS cells (Helbling et al., 2019) and expressed both NG2 and CD34, consistent with previous descriptions of periarteriolar and endosteal progenitors with restricted adipose potential (Morikawa et al., 2009; Kunisaki et al., 2013; Abdallah et al., 2015) (**Extended Data Fig. 7 and 8a**). However, contrary to a previous report suggesting that Gremlin-1 (*Grem1*) identifies a progenitor population with bone and cartilage potential (Worthley et al., 2015), we found poor expression of *Grem1* in MSC-Ss, with the published transcriptome of Grem1^+^ cells correlating more closely with downstream MPrs, in which expression of the osteoblast transcription factor Osterix (Osx) was also observed using *Osterix-Gfp* reporter mice (Rodda et al., 2006) (**Extended Data Fig. 7 and 8a,c**). Collectively, these data combined with our functional assays suggest two distinct sources of bone-forming mesenchymal cells in different spatial compartments of the BM niche, which can be resolved and prospectively isolated based on differential Sca-1 and LepR expression.

**Figure 4.**
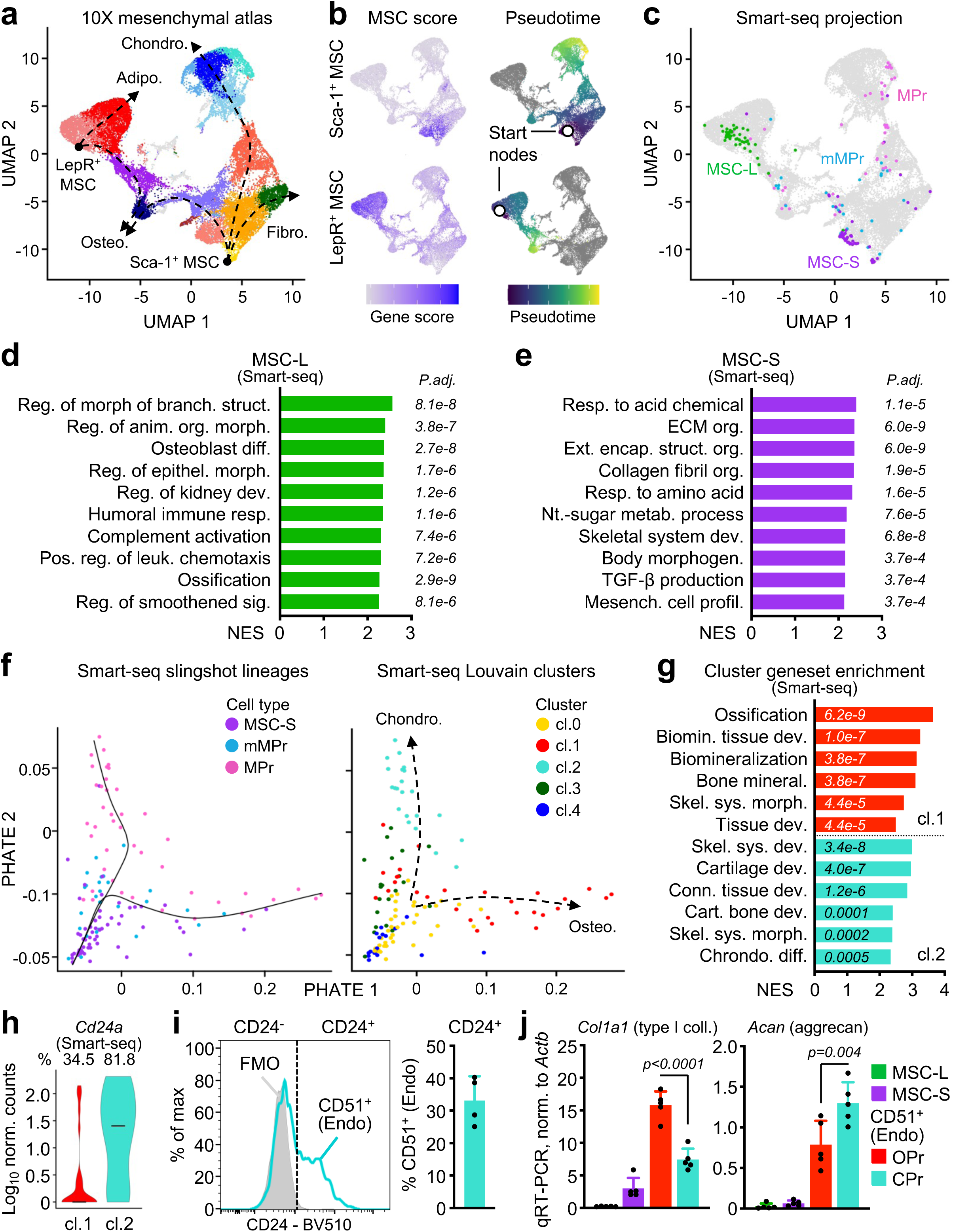
Mesenchymal cells have two distinct origin in the BM niche. **a,** UMAP of a 10X mesenchymal atlas created by integrating n = 3 published 10X scRNA-seq datasets of BM mesenchymal cells, with major lineage branches indicated. Chrondo.: chondroblastic, Adipo.: adipocytic, Osteo.: osteoblastic, Fibro.: fibroblastic. **b,** Expression of gene signatures associated with Sca-1^+^ or LepR^+^ MSCs (left) and pseudotime calculated with Monocle3 using start nodes corresponding to the highest MSC gene scores (right). **c,** Projection of Smart-seq scRNA-seq transcriptomes onto the 10X mesenchymal atlas. **d-e,** GSEA of selectively enriched GO biological pathways in MSC-L (d) and MSC-S (e). **f-h,** Assessment of adipogenic vs. chondrogenic differentiation: (f) Diffusion map created with PHATE from Smart-seq scRNA-seq datasets showing lineage pathways inferred by Slingshot (left) and Louvain clustering (right); and (g) GSEA of selectively enriched GO biological pathways (NES: normalized enrichment score, P.adj.: adjusted *P values*) and (h) violin plot showing expression of *Cd24a* and percentage of cells in which expression was detectable in cluster 1 (cl.1) and cluster 2 (cl.2) cells. **i,** Representative flow cytometry plots (left) and quantification (right) of the proportion of cells expressing CD24 in the CD51^+^ Endo fraction. FMO: fluorescence minus one. **j,** Expression of indicated genes measured by qRT-PCR in the indicated stroma populations isolated by flow cytometry. OPr: CD24^-^ subset of the CD51^+^ Endo fraction, CPr: CD24^+^ subset of the CD51^+^ Endo fraction. Results are normalized to expression of *Actb*. Data in (i) and (j) are means ± S.D. with points showing values for individual mice; *P. values*, one-way ANOVA with Tukey *post-hoc* test. For (d), (e), and (g), *P. values* are derived from permutation tests.

Projection of our Smart-seq populations onto the 10X mesenchymal atlas also revealed that individual (m)MPr cells were distributed across clusters containing osteoblast and chondroblast progenitors (**Fig. 4c**). To understand how cells deriving from MSC-Ss differentiate into these two lineages, we produced a diffusion map (Moon et al., 2019) of MSC-S/mMPr/MPr cells, and unbiased Slingshot analysis (Street et al., 2018) then identified two distinct lineages originating from MSC-Ss (**Fig. 4f**). Comparison of gene expression between these lineages revealed a branch significantly enriched for bone formation (cl.1) and another for cartilage development (cl.2) (**Fig. 4g**). Accordingly, gene expression of cells in cl.1 was highly correlated with cells expressing the committed osteoblast markers Osx and osteocalcin (Ocn) (Yu et al., 2015; He et al., 2017), whereas those in cl.2 were most correlated with primary chondrocytes (Mamidi et al., 2021) (**Extended Data Fig. 8d**). To determine whether cells in each lineage branch could be distinguished prospectively, we searched among DEGs for cell surface markers and identified expression of the mature chondrocyte marker *Cd24a* in a higher proportion of cartilage-biased cl.2 progenitors (**Fig. 4h**, **Extended Data Fig. 8e**) (Belluoccio et al, 2010). Using flow cytometry, we confirmed expression of CD24 in a subset of (m)MPr cells, with CD24^+^/CD51^+^ cells having significantly greater expression of the cartilage-specific protein aggrecan (*Acan*) and chondroblast lineage transcription factor *Sox9*, and thus representing CPrs (**Fig. 4i,j**; **Extended Data Fig. 8f,g**). Conversely, CD24^-^/CD51^+^ (m)MPr had higher expression of bone-associated type 1 collagen (*Col1a1*) and defining osteoblast transcription factor *Runx2*, thus representing OPrs. Collectively, these molecular data indicate that MSC-Ss differentiate into either osteoblasts or chondroblasts, with surface expression of CD24 enriching for progenitors biased toward cartilage development.

Finally, to explore how flow-defined stromal cells were engaging in interactions with HSPCs, we used the interactome package CellChat (Jin et al., 2021) to predict important receptor-ligand interactions between stromal populations from our 10X stroma map and oligo hashtag-labelled HSPCs analyzed by 10X scRNA-seq (**Extended Data Fig. 9a**). We found that MSC-Ls from both CM and Endo locations were the primary source of outgoing signals in the BM niche across many different classes of ligands sensed by both HSPCs and other stromal populations, highlighting the role of MSC-Ls as professional support cells for HSPCs as well as coordinators of both stromal compartments (**Extended Data Fig. 9b**). Conversely, SECs were the principal receivers of signals emanating from both stroma and HSPCs, consistent with their intimate connections with HSPCs and MSC-Ls in the same CM niche (**Extended Data Fig. 9b**). Collectively, these results uncover the complex interactions occurring within and between spatial compartments of the BM, which may be perturbed under disease conditions.

### Disruptions of the BM niche remodel various stromal compartments

Having optimized and validated a method for systematic profiling of niche compartments, we next asked how stromal cell populations were affected by stimuli that disrupt the BM niche and its resident hematopoietic cells. Acute myeloid leukemia (AML) is a common blood cancer that can be modeled effectively using HSPCs retrovirally transformed with the MLL/AF9 fusion protein to produce malignant blasts (Krivtsov et al., 2006). As previously reported (Galan-Diez et al., 2022), AML developed two-to-four weeks after injection of 2×10^5^ dsRed^+^ MLL/AF9 blasts in unconditioned WT C57Bl/6 mice (**Fig. 5a,b**), which was associated with global remodeling of both niche compartments, with increased numbers of all major stromal cell populations (**Fig. 5c; Extended Data Fig. 9c**). These results expand previous studies separately reporting increased numbers of SECs and LepR^+^ MSCs under similar conditions (Duarte et al., 2018; Xiao et al., 2018) by demonstrating that AML infiltration remodels the entire BM niche to its own advantage. This contrast with chronic myelogenous leukemia (CML) produced by inducible expression of the double *Scl-tTA::TRE-BCR/ABL* transgene that caused a specific expansion of (m)MPr cells to support tumor growth (Schepers et al., 2013), and with aging, where the same endosteal populations were depleted and adopted a pro-inflammatory phenotype marked by production of IL-1β that contributed to age-related HSPC dysfunction (Mitchell et al., 2023). These findings highlight the disparities in response of the BM niche to various injuries.

**Figure 5.**
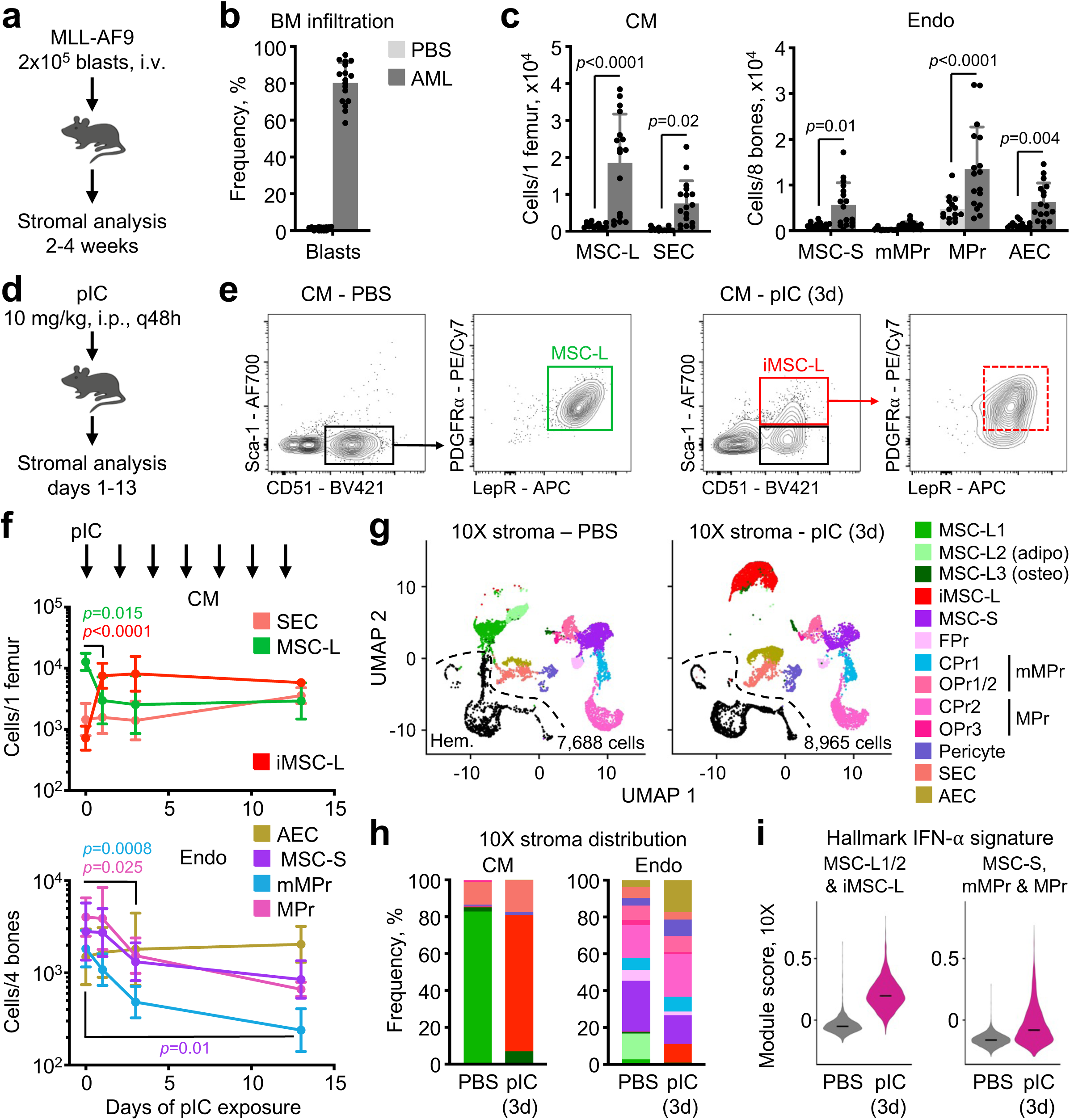
Neoplastic and inflammatory challenges remodel distinct niche compartments. **a-c**, Niche remodeling following acute myeloid leukemia (AML) development: (a) AML model produced by injecting retrovirally transformed dsRed^+^/MLL-AF9 blast cells into unconditioned mice; and (b) frequency of AML blasts among total BM cells and (c) quantification of indicated CM and Endo stromal cell populations 2-4 weeks after injection of AML blast cells. **d-f,** Niche remodeling following interferon-mediated inflammatory challenge: (d) inflammation model produced by repeated injection of polyinosinic:polycytidylic acid (pIC) every second day for up to 13 days; (e) representative flow cytometry plots showing increased expression of Sca-1 in inflammatory MSC-Ls (iMSC-L) in the CM of 3 day-pIC-injected mice; and (f) quantification of indicated CM and Endo stromal cell populations at indicated times following repeated pIC injections. **g-i,** 10X scRNA-seq profiling of CM and Endo fractions isolated from control (n = 1, 2 male mice) and 3 day-pIC-injected (n = 2, 2 male mice each) mice: (g) UMAP of merged datasets with number of analyzed cells and hematopoietic contaminants (Hem.) shown in black; (h) frequency of identified CM and Endo cell types; and (i) gene score for Hallmark interferon (IFN)-α geneset in indicated cell types. Data in (c) and (f) are means ± S.D. with points showing values for individual mice; *P. values*, two-way ANOVA with Sidak’s *post hoc* test.

### Inflammation selectively remodels MSC-Ls

To understand better how inflammation affects the BM niche, we serially administered the toll-like receptor 3 ligand polyinosinic:polycytidylic acid (pIC) every second day for up to 13 days (**Fig. 5d**), which induces type 1 interferon (IFN)-dependent inflammation and HSPC activation *in vivo* (Essers et al., 2009; Pietras et al., 2014). Remarkably, pIC caused a striking upregulation of Sca-1 on MSC-Ls to produce inflammatory MSC-Ls (iMSC-L) and led over time to a decline in all Endo mesenchymal populations (**Fig. 5e,f; Supplementary Fig. 3a**). Inflammatory MSC-Ls were detectable by flow cytometry within 3 days of pIC treatment (**Fig. 5f**), and 10X scRNA-seq analyses of niche compartments isolated from 3 day-pIC-exposed mice revealed clustering of iMSC-Ls away from steady state MSC-Ls, demonstrating dramatic divergence in their transcriptomic landscape and not just altered expression of the *Ly6a*/Sca-1 marker (**Fig. 5g; Supplementary Fig. 3b**). Identification of major stromal populations by scRNA-seq confirmed that the shift in MSC-L identity in both CM and Endo compartments was the greatest change observed upon pIC exposure, with less striking changes in frequency of all Endo mesenchymal subsets and AECs (**Fig. 5h**). The preferential response of MSC-Ls to pIC was associated with a greater upregulation of an IFN-α-regulated gene signature in CM MSC-L1/2/iMSC-L cells than in Endo MSC-S/(m)MPr cells (**Fig. 5i**). Consequently, we proposed that MSC-Ls were particularly sensitive to type 1 IFNs, possibly owing to greater resting expression of key components of the IFN signal transduction pathway (**Extended Data Fig. 9d**). To test this idea directly, we crossed mice carrying the IFN responsive *Mx1-Cre* promoter with a *Rosa26-mT/mG* reporter strain in which Cre recombinase converts tdTomato expression to GFP (Kuhn et al., 1995; Muzumdar et al., 2007). After pIC injections, we observed greater conversion to GFP expression in CM MSC-Ls than in any MSC-S-derived mesenchymal populations, confirming the heightened IFN sensitivity of MSC-Ls (**Extended Data Fig. 9e**). Collectively, these data demonstrate that inflammation and IFN signaling specifically affect CM MSC-Ls.

### Inflammatory MSC-Ls regulate monocyte dynamics by secreting chemokines

To understand the importance of iMSC-Ls in coordinated hematopoietic responses to pIC and IFNs, we isolated iMSC-L from 3 day-pIC-exposed mice for Smart-seq scRNA-seq analyses. By merging these data with Smart-seq analyses of control (Ctrl) MSC-Ls isolated from untreated mice and iMSC-Ls isolated from 24-month-old aged mice (**Fig. 6a; Supplementary Fig. 3c**) (Mitchell et al., 2023), we confirmed higher *Ly6a* and *Ly6e* (Sca-1) expression in pIC-exposed and aged iMSC-Ls (**Fig. 6b**). We also found marked changes in gene expression in pIC-exposed iMSC-Ls, with upregulation of multiple pathways related to IFN signaling and innate immune responses, and downregulation of pathways related to mesenchymal differentiation, including those linked to bone formation (**Fig. 6c,d**). Accordingly, pIC-exposed iMSC-Ls were no longer able to form CFU-F colonies *ex vivo* (**Fig. 6e**), which is a prerequisite for assays of bone forming capacity, whereas mixed Endo mesenchymal cells (MSC-S/(m)MPr) from 3 day-pIC-exposed mice had no impairment in colony formation (**Extended Data Fig. 10a**). Despite this, the trabecular bone mass of mice exposed to pIC over 13 days was unexpectedly increased (**Extended Data Fig. 10b,c**), which is probably attributable to the parallel role of pIC-induced IFN secretion in inhibiting osteoclast formation (Takayanagi et al., 2002).

**Figure 6.**
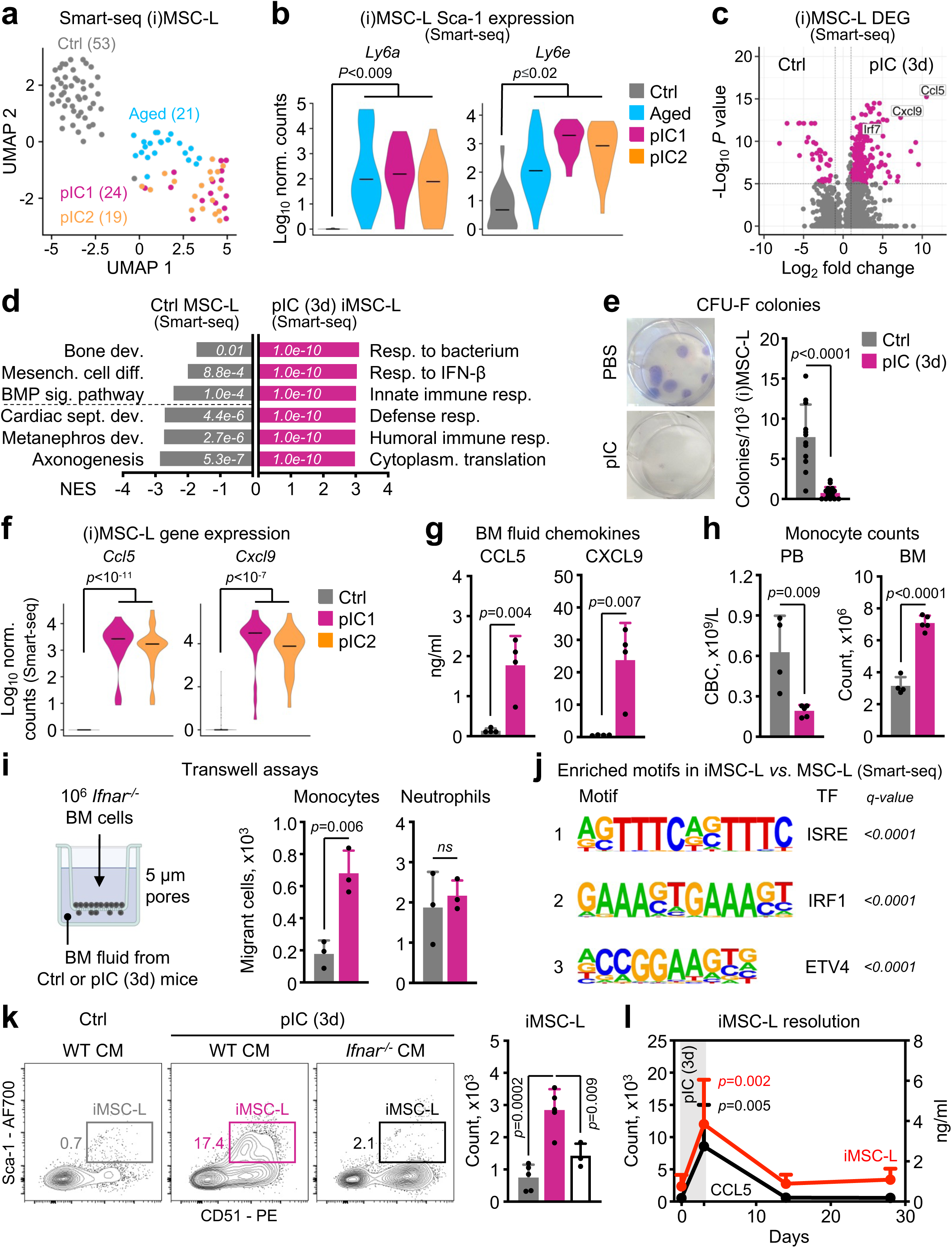
iMSC-Ls coordinate altered monocyte dynamics in the CM. **a-d,** Smart-seq scRNA-seq profiling of MSC-Ls isolated from young untreated control mice (Ctrl, n = 3 biological replicates, 2-3 male mice each) and iMSC-Ls isolated from 24-month-old mice (Aged, n = 1 male mouse) and 3 day-pIC-injected young mice (pIC1/2, n = 2 biological replicates, 1 male & 1 female mouse each): (a) UMAP of merged datasets with number of individual (i)MSC-L cell analyzed; (b) expression of genes encoding Sca-1 constituents; (c) volcano plot showing differentially expressed genes (red points show genes with log_2_ fold change >|1| and adjusted p value <10^-5^); and (d) GSEA of selectively enriched GO biological pathways (NES: normalized enrichment score, P.adj.: adjusted *P values*) in Smart-seq (i)MSC-Ls. **e,** Representative images (left) and quantification (right) of CFU-F obtained from CM (i)MSC-Ls isolated from 3 days PBS (Ctrl) or pIC-treated mice (1,000 cells/35-mm well cultured for 8 days in 5% O_2_/iROCK conditions). **f-g,** Changes in chemokine production: (f) expression of *Ccl5* and *Cxcl9* genes in indicated Smart-seq (i)MSC-Ls; and (g) production of CCL5 and CXCL9 in BM fluids of 3 days PBS (Ctrl) or pIC-treated mice. **h,** Monocytes counts in peripheral blood (PB) and BM of 3 days PBS or pIC-treated mice. **i,** Changes in monocyte migration with scheme of the transwell assays using 10^6^ c-Kit-depleted *Ifnar^-/-^* BM cells in the upper chamber and BM fluid from 3 days PBS (Ctrl) or pIC-treated mice in the bottom chamber (left), and quantification of the number of monocytes and neutrophils migrating into lower chamber after 2 hours (right). **j,** Top 3 transcription factor (TF) motifs enriched in genes differentially expressed in iMSC-Ls and identified using HOMER analysis. **k,** Representative flow cytometry plots (left) and quantification (right) of iMSC-Ls 3 days after injection of PBS (Ctrl) or pIC in either wild type (WT) or *Ifnar^-/-^*mice. **l,** Kinetics of iMSC-L resolution with line plots showing evolution in the numbers of CM iMSC-Ls and BM fluid concentration of CCL5 following an initial 3 days of pIC treatment (n = 4-5 mice per time point). Data in (b) and (f) are violin plots of Log10 normalized (norm.) Smart-seq counts; *P. values,* Wilcoxon rank sum test. Data in (e), (g), (h), (i), (k), and (l) are means ± S.D. with points showing values for individual mice; *P. values* for (e), (g), (h), and (i), Student’s t test; *P. values* for (k) and (l), one-way ANOVA with Tukey’s *post-hoc* test. *P. values* for (d) are derived from permutation tests.

Among the genes significantly upregulated in pIC-exposed iMSC-Ls were multiple chemokines, particularly CCL5 (RANTES) and CXCL9 (MIG) (**Fig. 6f**). We confirmed increased concentrations of both chemokines in BM fluids of 3 day-pIC-exposed mice (**Fig. 6g**). Given the known effect of CCL5 in promoting myeloid bias in HSCs (Ergen et al., 2012), we next asked whether this local production of inflammatory chemokines in the BM niche could be important in the hematopoietic response to pIC. To test this notion, we conducted CellChat analysis to explore putative interactions between stromal cells and mature BM cells isolated from control and 3 day-pIC-exposed mice (**Extended Data Fig. 10d,e**). This analysis indicated a replacement of hematopoietic cells by iMSC-Ls as the predominant producers of CCL chemokines, and predicted that neutrophils and monocytes would be the principal receivers of these signals (**Extended Data Fig. 10f,g**). Interestingly, we found a marked redistribution of mature CD11b^+^/Ly6C^hi^ monocytes upon pIC treatment, with relative depletion in peripheral blood (PB) and expansion in BM (**Fig. 6h; Supplementary Fig. 3d**), whereas neutrophils were depleted in the BM over the same time period (**Extended Data Fig. 10h**). To determine whether local chemokine production by iMSC-Ls was responsible for the selective retention of monocytes in the BM, we extracted BM fluid from control and 3 day-pIC-exposed mice to evaluate its chemoattractant activity in transwell assays (**Fig. 6i**). Importantly, we used BM cells lacking the type I IFN receptor (*Ifnar^-/-^*) (Prigge et al., 2015) to remove any confounding effect of IFNs present in the BM fluid of pIC-exposed mice on migrating hematopoietic cells. We found that BM fluid from pIC-injected mice specifically attracted monocytes, but not neutrophils, across the transwell membrane to a greater extent than BM fluid from control mice (**Fig. 6i**), confirming a role for altered MSC-L activity in abnormal monocyte dynamics. To explore the relevance of this finding, we compared gene expression in monocytes from control and 3 day-pIC-injected mice, finding that pIC exposure increased expression of markers of activation like cathepsin G and MHC class I (**Extended Data Fig. 10i**), which we confirmed by flow cytometry (**Extended Data Fig. 10j**). Additionally, we found that monocytes from pIC-injected mice had significantly greater reactive oxygen species (ROS) content (**Extended Data Fig. 10k**), which has been associated with HSC activation and enhanced myelopoiesis in previous studies (Zhu et al, 2017; Mistry et al, 2019). Together, these results show that pIC-induced production of chemokines by inflammatory MSC-Ls that attracts and/or retains tissue-toxic monocytes in the BM niche, which might also contribute to IFN-mediated HSPC activation and/or functional impairment.

To understand how MSC-Ls are reprogrammed by pIC, we investigated the TF motifs that were enriched among genes differentially regulated in iMSC-Ls using HOMER. We found striking enrichment for interferon-stimulated response element (ISRE) motifs, as well as motifs associated with interferon-response factors (IRF) (**Fig. 6j**), with the gene encoding IRF7 being particularly upregulated in pIC-exposed iMSC-Ls (**Extended Data Fig. 10l**). Accordingly, we did not observe Sca-1 upregulation or iMSC-L emergence upon injection of pIC in *Ifnar^-/-^* mice (**Fig. 6k**). We also found that the stromal response to IFN dissipated upon pIC withdrawal, with iMSC-Ls no longer observed in the CM and CCL5 levels returning to baseline 2 weeks after the last pIC injection (**Fig. 6l**). Collectively, our data suggest that the enhanced IFN sensitivity of MSC-Ls poises them to act as critical first responders in the BM niche upon exposure to pIC and IFNs, transducing these pathogenic signals through production of inflammatory chemokines that modulate the number and activity of neighboring cells in the CM and directly affect the trafficking of inflammatory monocytes in and out of the marrow.

## Discussion

Assessing how stromal cells respond to stimulation is necessary to understand how diverse challenges, such as cancer and inflammation, remodel the BM niche to affect HSPC activities. Here, we present a unified scheme for stromal cell identification and isolation, and demonstrate its use to profile BM niche changes in the context of different perturbations. Through extensive validation and cross-referencing to previous datasets, we find that our annotation captures most major cell types described in the BM niche and aligns well with previous imaging and functional studies. In particular, we corroborate the existence of two distinct vascular niches and the key role of MSC-Ls as main interactants with HSPCs and other stromal cells. Importantly, many of our findings have recently been replicated in human BM, with a population of Fibro-MSCs enriched in the endosteal region that closely resembles mouse MSC-Ss, and populations of THY1^+^ and Adipo-MSCs that are similar to mouse MSC-Ls (Bandyopadhyay et al., 2024), illustrating the utility of our approach in capturing similar cell types and spatial partitions to those observed in human patient samples. Moreover, our functional assays and analysis of previously published sequencing studies sheds new light on the developmental relationships of mesenchymal cells, positing the existence of two distinct but converging sources of bone-forming cells from either CM MSC-Ls or Endo MSC-Ss, which partially overlaps with recent findings on the roles of different skeletal stem cells in fracture repair (Jeffery et al, 2022). We also find that fibroblast and chondroblast potential is restricted to MSC-Ss and their derivatives and that MSC-Ls alone are predicted to generate adipocytes *in vivo*, even though both MSC-Ss and MSC-Ls can be induced to form CFU-F and undergo adipogenic differentiation *ex vivo*. Since lineage tracing studies in mice show that MSC-L contribution to bone is first detectable at 2 months of age and then increases progressively with aging (Zhou et al., 2014), we speculate that MSC-Ss and their derivatives could have a more important role in early bone development and extension, which is also consistent with their greater capacity for chondrocyte formation needed at the growth plate. Our work further implies that different mesenchymal lineage fates are largely partitioned according to spatial location in the BM niche under steady state conditions in adult mice, though others have shown that this lineage restriction may be negotiable under stress conditions, such as when MSC-Ls produce chondroblasts during fracture repair (Zhou et al, 2014), or when MSC-Ss produce occasional adipocytes upon transplantation (Morikawa et al., 2009). Remaining questions to be investigated include whether different mesenchymal lineages are induced or maintained in different spatial niches by extrinsic signals or by cell-intrinsic differentiation programs, and whether different MSCs have distinct developmental origins.

Our analysis of stromal cells under stimulation conditions demonstrates that spatial compartmentalization is an important determinant of the response to specific perturbations. In particular, we find the CM to be exquisitely sensitive to pIC exposure, showing marked changes in stromal cell abundance and/or activity. Previous studies demonstrated that MSC-Ls are responsive to pIC and viral infections (Helbling et al, 2019; Isringhausen et al, 2021), and we build upon those works by showing that iMSC-L forego mesenchymal differentiation in favor of local production of inflammatory chemokines, which modulate monocyte dynamics in and out of the marrow. MSC-Ls are already known to regulate monocyte egress from the BM niche through production of chemokines like CCL2 (Shi et al., 2011), and we show here how this chemokine landscape can be specifically altered by the type I IFN response to affect monocyte balance during inflammation. The retention of monocytes with a pro-inflammatory phenotype and active respiratory burst is also likely to directly contribute to the hematopoietic response to pIC, as supported by studies reporting modulation of myeloid differentiation by extracellular ROS acting directly on GMPs (Zhu et al., 2017). Moreover, we show that MSC-Ls have enhanced capacity to respond to IFNs, which did not extend to endosteal mesenchymal cells and explains the specificity of pIC effects. This finding complements our recent work, in which we observed a similar population of iMSC-Ls in aged mice (Mitchell et al, 2023), and suggests that repeated or chronic low-level exposure to IFNs in the aging niche could be implicated in the expansion and functional impairment of aged MSC-Ls to the presumed detriment of the HSPCs with which they interact. Moreover, in practical terms, we confirm that MSC-Ls are the stromal cell population mainly affected by *Mx1*-Cre, which is an important confounding factor in the use of this model system for studying HSC functions (Park et al., 2012; Velasco-Hernandez et al., 2016).

Collectively, our work reconciles numerous studies of the BM niche into a unified scheme that incorporates its inherent spatial compartmentalization and provides a method for investigating the effect of perturbations on both hematopoietic and stromal compartments. Further studies can now be directed at understanding how this stromal niche compartmentalization is established and maintained, and how specific stromal responses might be manipulated therapeutically to modulate inimical hematopoietic responses in inflammation and cancer.

**Extended Data.** See accompanying document.

## Acknowledgements

We thank Drs. J. Butler (University of Florida), M. Maryanovich (Albert Einstein College of Medicine), L. Ding (Columbia University), S. Kousteni (Columbia University), and I. Aifantis (New York University) for sharing mouse lines and other reagents, M. Kissner for management of the CSCI Flow Cytometry Core facilities, and all members of the Passegué laboratory for critical insights and suggestions. J.W.S. was supported by EMBO postdoctoral fellowship ALTF-2021-196 and Damon Runyon Cancer Research Foundation DRG-2493-23 (William Raveis Family Fellowship), E.V.V. by a Rubicon Grant from The Netherlands Organization for Scientific Research, a Stem Cell Grant from BD Biosciences, and a NYSTEM training grant, and M.A.P. by NIH TL1DK136048. F.J.C-N., X.W. and B.G. were supported by grants from the Wellcome (206328/Z/17/Z), CRUK (C1163/A21762) and core funding by Wellcome to the Cambridge Stem Cell Institute. This work was funded by NIH R01CA184014 and NIH R35HL135763 to E.P. and supported in part through the NIH/NCI Cancer Center Support Grant P30CA013696 to CUIMC.

## Author contributions

Conceptualization, E.V.V., J.W.S., R.Z., and E.P.; methodology, E.V.V., J.W.S., R.Z., F.J.C.-N., X.W., P.T.S., E.G., B.G.; investigation, E.V.V., J.W.S., R.Z., M.A.P., F.J.C-N., X.W., and P.T.S.; visualization, E.V.V., J.W.S., and R.Z.; funding acquisition, B.G. and E.P.; project administration, E.P.; supervision, E.P.; writing – original draft, J.W.S., E.V.V., R.Z., and E.P.; writing – review & editing, J.W.S., R.Z., and E.P.

## Additional information

Supplementary information is available for this paper. Correspondence and requests for materials should be addressed to E.P.

## Methods

### Mice

All animal experiments were conducted at the University of California San Francisco (UCSF) or Columbia University Irving Medical Center (CUIMC) in accordance with Institutional Animal Care and Use Committee protocols approved at each institution, and in compliance with all relevant ethical regulations. Young wild type (WT) C57BL/6-CD45.2 mice were purchased from Jackson Laboratory and bred in house. Aged WT C57BL/6-CD45.2 mice were obtained from the National Institute on Aging (NIA) when they were 18 months old and used for experiments when they were 24 months old. BALB/CJ, *Ifnar1*^-/-^, *Mx1*-Cre, and mTmG reporter mice were purchased from the Jackson Laboratory. For mouse strains expressing fluorescent reporters, *Vegfr3*-YFP and *CX40*-GFP mice were shared from the laboratory of Dr. Jason Butler at the University of Florida. *Nestin*-GFP mice were shared from the laboratory of Dr. Maria Maryanovich at Albert Einstein College of Medicine. *Lepr*-Cre:ROSA26-YFP and *Prx1*-Cre:ROSA26-tdTomato mice were shared from the laboratory of Dr. Lei Ding at CUIMC. *Osterix*-GFP mice were shared from the laboratory of Dr. Stavroula Kousteni at CUIMC. *Hes1*-GFP mice were shared from the laboratory of Dr. Ioannis Aifantis at New York University. Mice were 8 to 12 weeks of age when used for experiments. No specific randomization or blinding protocol was used with respect to the identity of experimental animals, and both male and female animals were used in all experiments.

### In vivo assays

For the MLL-AF9 model, BM cells from leukemic mice injected with dsRed^+^/MLL-AF9 blasts were kindly provided by the laboratory of Dr. Stavroula Kousteni at CUIMC and mice were then injected intravenously into the tail vein (2×10^5^ cell/mouse), with BM and stroma analyses completed 3 to 4 weeks later. For polyinosinic:polycytidylic acid (pIC, Cytiva) treatment, pIC was dissolved in sterile PBS at 1.25 mg/ml and then mice were injected intraperitoneally with 10 mg/kg pIC every 48h, with BM and stroma analyses performed on days 1-13 after starting pIC treatment. For dragon green bead (DGB, Bangs Laboratories, FSDG001) analyses, mice were injected retro-orbitally with 2.5 μl/g DGB solution 10 minutes before euthanasia and then perfused with 20 ml PBS by cardiac puncture before collecting the bones for analysis.

### Combined stromal and hematopoietic cells isolation

Mice were euthanized in a rising concentration of carbon dioxide, followed by cervical dislocation. For each mouse, both femurs, tibiae, hemipelves, and humeri were dissected and thoroughly cleaned using KimWipes (Kimtech) and were used for combined isolation of stromal and hematopoietic cells. To isolate central marrow (CM) stromal cells, the proximal and distal epiphyses were removed from 1 femur. Intact marrow plugs were flushed with Hank’s balanced saline solution (HBSS) without calcium or magnesium into 5 ml polypropylene tubes by inserting a 3 ml syringe with 22G needle into the distal end of the femur. The plugs were digested with 1 ml of a solution of 3 mg/ml type I collagenase (Worthington) dissolved in HBSS for 10 min at 37 °C with 110 rpm shaking. The tube was then vortexed briefly and the supernatant was removed into a new tube through a 100 μm mesh, taking care not to disturb the marrow plug. A further 1 ml of 3 mg/ml type I collagenase solution was then added to the marrow plug and the digestion repeated. After the second incubation, the plug was dissociated by pipetting up and down with a P1000 pipette before passing the cell suspension through a 100 μm mesh into the tube containing the previous digestion supernatant. To isolate endosteal (Endo) stromal cells and hematopoietic cells, the flushed femur was combined with the other bones, gently crushed up to 10 times using a mortar and pestle, and thoroughly washed with 10 ml HBSS until all the non-adherent BM cells were removed and collected in a separate 15 ml polypropylene tube. The bone chips were then digested in a 15 ml polypropylene tube with 3 ml of 3 mg/ml type I collagenase solution for 1 hour at 37 °C with 110 rpm shaking. After digestion, the tube was vortexed briefly and the cell suspension was filtered through a 100 μm mesh into a new tube. The bone chips were then washed with HBSS, and the washing was collected into the same tube as the digestate. For cell suspensions acquired for both stromal compartments, red blood cells were removed by adding 1 ml ACK lysis buffer (150 mM NH_4_Cl and 10 mM KHCO_3_) and incubated on ice for 3 minutes before washing with HBSS containing 4% fetal bovine serum (HI FBS, Gibco). During the optimization of the protocol outlined above, the digestion with 3 mg/mL type I collagenase was performed in parallel to and compared with 3 mg/mL type I collagenase + 4 mg/mL type II Dispase (Roche), 250 μg/mL Liberase DL (Roche) + 200 U/mL DNAse I (Sigma), and mechanical dissociation with P1000 pipette. The BM cells collected from crushed bones were resuspended in HBSS with 2% FBS and RBCs were removed by lysis with ACK buffer. The BM cell suspension was further purified on a density gradient (Histopaque 1119, Sigma-Aldrich), by layering 2 ml of the cell suspension under 2 ml of Histopaque solution. For profiling of CM plug and Endo marrow, only femurs were used and were flushed once with a 3 ml syringe and 21G needle to collect BM plugs, while the flushed bones were crushed to collect the remaining Endo-associated BM cells. No ACK RBC lysis was performed for these analysis. Both stromal and hematopoietic cells were finally counted using a Vicell automated cell counter (Beckman Coulter).

### Flow cytometry of stromal cells

For stromal cell profiling, Endo and CM cell preparations were stained with CD45-APC/Cy7 (BD, 557659; 1:400), Ter119-PE/Cy5 (Invitrogen, 15-5921-83; 1:400), Sca-1-AF700 (eBioscience, 56-5981-82; 1:800), CD31-PE (BD, 553373; 1:200), CD105-BV786 (BD, 564746; 1:200), CD51-BV421 (BD, 740062; 1:100), LepR-biotin (R&D, BAF497; 1:100), and PDGFRα-PE/Cy7 (Invitrogen, 25-1401-82; 1:100) for 30 minutes on ice. Cells were then washed and stained with Streptavidin-APC (BioLegend, 405207;1:400) for 15 minutes on ice. For analysis of CD24 expression, CD24-BV510 (BioLegend, 101831; 1:400) was added to the staining panel. For the analysis of endomucin, endomucin-APC (Invitrogen, 50-5851-80; 1:100) was used and LepR antibody was dropped from the staining panel. For the analysis of CD34, CD34-FITC (eBioscience, 11-0341-85; 1:25) was added to the staining panel. For the analysis of NG2, cells were incubated with unconjugated monoclonal rabbit anti-mouse NG2 antibody (Millipore Sigma) alongside other conjugated antibodies, then washed and stained with goat anti-rabbit secondary antibody conjugated to AF488 (Invitrogen, 1:400) for 30 minutes. Cells were finally washed and resuspended in HBSS with 4% FBS and 1 μg/ml propidium iodide and then analyzed or sorted on purity mode on a BD FACSAria II (UCSF) or FACSAria II SORP (CUIMC). Data collection was performed using FACSDiva (v.9) and analysis was performed using FlowJo (v.9/v.10).

### Flow cytometry of hematopoietic cells

For HSPC profiling, BM cells were stained with c-Kit-APC-Cy7 (BioLegend, 105826; 1:800), Sca-1-BV421(BioLegend, 108128; 1:400), CD150-BV650 (BioLegend, 115931; 1:200), CD48-A700 (BioLegend, 103426; 1:400), Flt3-PE (eBioscience,12-1351-82; 1:100), CD34-FITC (eBioscience, 11-0341-85; 1:25) and CD16/32-PE/Cy7 (BioLegend, 101318; 1:800), along with lineage antibodies in PE/Cy5 (Gr1 [eBioscience, 15-5931-82; 1:800], CD11b [Invitrogen, 15-0112-82; 1:800], B220 [Invitrogen, 15-0452-82; 1:800], CD19 [BioLegend, 115510; 1:800], CD5 [BioLegend, 100610; 1:800], CD4 [Invitrogen, 15-0041-82; 1:800], CD8a [Invitrogen, 15-0081-82], Ter119 [Invitrogen, 15-5921-83; 1:400], CD3 [Invitrogen, 15-0031-83; 1:400]). For mature cell profiling, cells were stained with Ly6G-FITC (BioLegend, 127606, 1:400), Ly6C-APC (BioLegend, 128016, 1:400), CD8-AF700 (BD, 557959, 1:400), B220-APC/Cy7 (BioLegend, 103224, 1:800), CD4-BV510 (BioLegend, 100553, 1:400), CD11b-BV786 (BioLegend, 101243, 1:800), CD115-PE (BioLegend, 135506, 1:100), and CD3-PE/Cy7 (BioLegend, 100220, 1:100). For some analyses, cells were also stained with H2-Kb-FITC (MHC class I, BioLegend, 116505, 1:100) or CD16/32-PE/Cy7 (BioLegend, 101318, 1:800). For DHR123 staining, after surface staining, cells were re-suspended in DHR123 (Invitrogen, D23806) at 1 μM in HBSS and incubated for 30 minutes at 37°C. Cells were finally washed, resuspended in HBSS with 2% FBS and 1 μg/ml propidium iodide, and analyzed on an Agilent Novocyte Quanteon or Penteon (CUIMC). Data analysis was performed using FlowJo (v.9/v.10)..

### Stromal and hematopoietic cell isolation for 10X scRNA-seq analyses

For sequencing of Endo and CM stromal cell preparations, 30,000-50,000 Ter119^-^/CD45^-^ cells were sorted into 1.5 ml tubes containing 300 μl of 50% FBS and 50% HBSS. For sequencing of BM cells, 40,000 live (propidium iodide negative) cells were similarly isolated from PBS or pIC-injected mice. For sequencing of defined HSPCs, 8,000 cells of 10 different populations (HSC, stHSC, MPP2, FcγR^-^ MPP3, FcγR^+^ MPP3, MPP4, GMP, MkP, CLP, CFU-E) were each sorted in separate 1.5 ml tubes and labelled with 0.05 μg of oligo-hashed antibodies specific for MHC class I and CD45 (BioLegend, TotalSeq B1-10) for 30 minutes on ice before being washed three times with HBSS containing 2% FBS, and then pooled together for subsequent partitioning. For sequencing of defined stromal populations, oligo-hashed antibodies at a concentration of 0.375 μg of antibody per 10^6^ cells were mixed with surface antibodies during stromal cell isolation from separate samples of CM or Endo preparations (1 TotalSeq B antibody per sample type). Cells were then washed three times with HBSS containing 4% FCS and the indicated stromal populations were isoled by flow cytometry and mixed according to the following scheme to produce two separate combined stains, each containing a total of 25-40,000 cells:

| Sample type | Oligo hash antibody | Combined stain 1 (cells) | Combined stain 2 (cells) |
| --- | --- | --- | --- |
| CM | TotalSeq B1 | MSC-L (6899) | SEC (2228) |
| CM | TotalSeq B2 | mMPPr (2775) | SEC (2097) |
| CM | TotalSeq B3 | mMPPr (2406) | SEC (2529) |
| CM | TotalSeq B4 | MPr (847) | SEC (1041) |
| CM | TotalSeq B5 | MPr (802) | SEC (1353) |
| CM | TotalSeq B6 | MSC-L (7642) | AEC (438) |
| CM | TotalSeq B7 | - | AEC (496) |
| Endo | TotalSeq B8 | MPr (10048) | MSC-S (8212) |
| Endo | TotalSeq B9 | AEC (5046) | MSC-L (2319) |
| Endo | TotalSeq B10 | SEC (3537) | mMPPr (6823) |

In all cases, following isolation, cells were rested for 1 hour on ice then pelleted at 350 x *g* for 5 minutes at 4°C before the supernatant was removed down to a volume of 40 μl. GEM generation and 3’ RNA library preparation was performed according to 10X Genomics protocol CG000315 Rev E, targeting 5000 cell data recovery. RNA libraries were pooled 1:1:1:etc, sequenced on an Illumina NovaSeq 5000, and aligned using Cellranger (v.7.0.1) to mouse genome mm10. For hashing samples, hashing libraries were pooled with RNA libraries at a ratio of 1:4. Library concentrations and fragment sizes were evaluated using Qubit dsDNA HS assay kit (ThermoFisher Scientific) and TapeStation D5000 DNA ScreenTape analysis (Agilent).

### Stromal cells and MSC-L isolation from Smart-seq scRNA-seq analyses

For Smart-seq analyses of Endo (AEC, MSC-S, mMPr, MPr) and CM (SEC, MSC-L) stromal cells isolated from young WT mice, we utilized our own published data (Mitchell et al., 2023) that was processed using the same analytical pipeline. For Smart-seq of iMSC-L isolated from 3 day-pIC-injected young WT mice and 24-month-old aged WT mice, we utilized the SMART-Seq Single Cell PLUS kit (96 reactions, Takara Bio). Single iMSC-L were sorted into 96 well plates containing 12.5 μl of CDS sorting solution provided in the kit. Preparation and amplification of cDNA and single cell libraries were completed according to the manufacturer’s instructions. Single cell libraries that passed quality control (n=43 from pIC-injected mice and n=30 from aged mice) were pooled at a 1:1:1:etc ratio before sequencing on an Illumina NextSeq 500/550 instrument using a version 2 kit (Illumina). Fastq files were generated for each library in Illumina BaseSpace, and quality of data was evaluated with FastQC and MultiQC (Andrews, 2010). After removing samples with poor sequencing quality, 43 cells from pIC-injected mice and 21 cells from aged mice were used for analysis. After trimming adapters with TrimGalore (Krueger, 2024), sequencies were pseudoaligned to the murine mm10 genome and counted using Salmon (Patro et al., 2017).

### 10X scRNA-seq data analyses - CM and Endo preparations

Count matrices for 10X scRNA-seq were analyzed with Seurat (versions 4 and 5) (Hao et al., 2021; Hao et al., 2024) in R (version 4.3.1). For CM and Endo CD45^-^/Ter119^-^ stromal samples, we observed higher counts of ambient RNA in some samples, so we adjusted the matrices for all stromal samples using the R package SoupX (Young et al., 2020) with default parameters. To exclude damaged cells or probable doublets, we filtered cells according to the following criteria after creation of Seurat objects: mitochondrial reads per cell >5.5%, number of reads per cell >65,000, number of unique genes per cell <1000 and >7000. For generation of the reference 10X stroma map, we merged the CM and Endo sequencing results of three independent datasets, normalized the data using the *SCTransform* function, and then integrated these samples together using SCT integration functions (*SelectIntegrationFeatures*, *PrepSCTIntegration*, *FindIntegrationAnchors*, *IntegrateData*). We performed principal component analysis (*RunPCA*, 35 components) before creating UMAP visualization (*RunUMAP*, 17 dimensions) and discovering Louvain clusters (*FindNeighbors*, 30 dimensions, *FindClusters*, resolution 0.5). We removed a single cluster that contained cells overlapping with numerous other clusters in the UMAP visualization, which probably represented a small number of residual cells of poor quality. We mapped Smart-seq dataset of stromal cells and 10X scRNA-seq of purified stromal populations to this 10X stroma reference map using *FindTransferAnchors* and *MapQuery* functions. We annotated clusters and assigned names manually in the scRNA-seq reference dataset by (i) evaluating key marker genes identified using the *FindAllMarkers* function, (ii) mapping our Smart-seq dataset, and (iii) mapping our purified 10X scRNA-seq samples. We evaluated correlations between clusters in the reference 10X stroma map and Smart-seq datasets by generating pseudobulk RNA profiles for each Smart-seq population (using the *AggregateExpression* function) then using the *clustify* function from the ClustifyR package (Fu et al., 2020) to calculate spearman correlations for each cluster in the reference 10X stroma map with the top 2,000 highly variable genes. For heatmap visualization, we scaled these correlations between 0 and 1 for each Smart-seq cell type. For analysis of the impact of pIC on stromal cells, we merged the sequencing results from control and 3 day-pIC-exposed mice to generate a complete dataset, before applying the same quality control filters and analytical functions as described above. To evaluate changes in gene expression within clusters upon pIC exposure, we used the *FindMarkers* function (minlog2foldchange 0.25, min.pct 0.25). To evaluate the type I IFN response in stromal cells, we scored cells in the merged dataset using the *AddModuleScore* function, utilizing the Hallmark Interferon Alpha Response v.7.5.1 signature. Geneset enrichment analysis was performed using the GO biological pathways with the *gseGO* function from clusterProfiler (Yu et al., 2012; Wu et al., 2021) (OrgDb = org.Mm.eg.db, ont = “BP”, minGSSize = 10, maxGSSize = 800, pvalueCutoff = 0.05, pAdjustMethod = “none”).

### 10X scRNA-seq data analyses - BM cells

Count matrices for 10X scRNA-seq were analyzed with Seurat (versions 4 and 5) in R (version 4.3.1). To exclude damaged cells or probable doublets, we filtered cells according to the following criteria after creation of Seurat objects: mitochondrial reads per cell >10%, number of unique genes per cell <200. To evaluate the effect of pIC on mature BM cells, we normalized control and pIC datasets using the *SCTransform* function, then integrated these samples together using SCT integration functions (*SelectIntegrationFeatures*, *PrepSCTIntegration*, *FindIntegrationAnchors*, *IntegrateData*). We scored the cell cycle status of these cells (*CellCycleScoring*) before regressing the effects of the S and G2M scores. We performed principal component analysis (*RunPCA*, 30 components) before creating UMAP visualization (*RunUMAP*, 30 dimensions) and discovering Louvain clusters (*FindNeighbors*, 30 dimensions, *FindClusters*, resolution 0.5). We annotated clusters using characteristic marker genes found with the *FindAllMarkers* function, and we evaluated the effect of pIC within clusters using the *FindMarkers* function (minlog2foldchange 0.25, min.pct 0.25).

### 10X scRNA-seq data analyses - Oligo-hashed stromal cells and HSPCs

Count matrices for 10X scRNA-seq were analyzed with Seurat (versions 4 and 5) in R (version 4.3.1). The RNA data for oligo-hashed stromal cells and HSPCs were filtered and analyzed as described in the preceding sections. Hashing data for HSPC and stromal datasets were normalized using *NormalizeData* then deconvoluted using *MULTIseqDemux* to assign labels for each cell, and merged into a single Seurat object.. For oligo-hashed stromal cells, since the frequency of successful hash labels was rather low across the 2 combined stains we generated, we utilized a combination of information to identify CM and Endo MSC-Ls, SECs, and AECs, including: any available hashtag labels, the presence of a cell type in each sample compared to the expected composition, and the presence of characteristic gene markers identified with the *FindMarkers* function. Data from the purified stromal samples were mapped to reference 10X stroma map using *FindTransferAnchors* and *MapQuery* function.

### 10X scRNA-seq data analyses – CellChat interactome

For simplicity, FcγR^+^ MPP3 were removed from the HSPC dataset and only cells corresponding to major stromal populations (MSC-L1, MSC-L2, MSC-S, mMPr, MPr, SEC, AEC) were extracted from the reference 10X stroma map. Raw counts from hematopoietic and stromal cells were then merged into a single Seurat object. Counts were normalized with *SCTransform*, and the normalized matrix was extracted to create a CellChat object (Jin et al., 2021) using *createCellChat*, which was then processed using standard functions and default settings, selecting only “Secreted Signaling” for analysis. We used the function *netAnalysis_signalingRole_heatmap* to create heatmaps showing major incoming and outgoing signals for included cell types across significantly enriched signaling families. We performed similar analysis to create CellChat objects for mature hematopoietic cells from control mice (neutrophils, monocytes, dendritic cells, granulocyte macrophage progenitors, B cells, and T cells) merged with control stromal cells (MSC-L1/2, MSC-S, mMPr, MPr, SEC, AEC), and from 3-day-pIC-exposed mice merged with pIC(3d) stromal cells (including iMSC-L instead of MSC-L1/2). We then used *mergeCellChat* to create a composite object, followed by *netAnalysis_signalingRole_network* and *netVisual_individual* for visualization of important pathways.

### Smart-seq scRNA-seq data analyses

The counts table derived from Mitchell et al., 2023 was imported into Seurat (versions 4 and 5) in R (version 4.3.1), with no additional filtering. Counts were processed using *NormalizeData*, *FindVariableFeatures*, and *ScaleData*. Differences in gene expression between stromal populations were evaluated using *FindMarkers* (minlog2foldchange 0.25, min.pct 0.25). For evaluation of differentiation trajectories among endosteal stromal cells, MPr, mMPr, and MSC-S cells were extracted to generate a diffusion map using the function *phate* in the R package PhateR (Moon et al., 2019). Dimension reduction and clustering were repeated on this dataset using the following functions and settings: *RunPCA* (30 dimensions), *RunUMAP* (20 dimensions), *FindNeighbors* (20 dimensions), *FindClusters* (resolution = 1.0). The function *slingshot* from the R package Slingshot (Street et al., 2018) was then used to identify trajectories in the PHATE dimension reduction, specifying cluster 4/MSC-S group as the starting node, and these trajectories were visualized using *getLineages* and *getCurves* functions before plotting in base R. Geneset enrichment analysis was performed using the GO biological pathways with the *gseGO* function from clusterProfiler (OrgDb = org.Mm.eg.db, ont = “BP”, minGSSize = 10, maxGSSize = 800, pvalueCutoff = 0.05, pAdjustMethod = “none”). For evaluation of the impact of aging and pIC on MSC-Ls, young control MSC-L data were downloaded from Mitchell et al. (2023), reprocessed, and merged with the pIC/aging iMSC-L Smart-seq counts generated in this study. These counts were then imported into Seurat (versions 4 and 5) in R and processed using *NormalizeData*, *FindVariableFeatures,* and *ScaleData*. We performed principal component analysis (*RunPCA*, 30 components) before creating UMAP visualization (*RunUMAP*, 20 dimensions) and discovering Louvain clusters (*FindNeighbors*, 20 dimensions, *FindClusters*, resolution 1.0). Differentially expressed genes were evaluated using *FindMarkers* (minlog2foldchange 0.25, min.pct 0.25), and volcano plots were generated using the *EnhancedVolcano* function from EnhancedVolcano. Geneset enrichment analysis was performed using the GO biological pathways with the *gseGO* function from clusterProfiler (OrgDb = org.Mm.eg.db, ont = “BP”, minGSSize = 10, maxGSSize = 800, pvalueCutoff = 0.05, pAdjustMethod = “none”). Evaluation of transcription factor motifs in differentially expressed genes was performed using HOMER (Heinz et al., 2010), running the *findMotifs.pl* function in the command line with default parameters for mouse promoters.

### Re-analyses of published 10X scRNA-seq datasets

To generate the 10X mesenchymal atlas, data were obtained from three previous studies: Baccin et al., 2020 [GSE122465]; Baryawno et al., 2019 [GSE128423]; Zhong et al., 2020 [GSE145477].

For GSE122465, the counts table and metadata (including principal components) were downloaded directly, with no further filtering. For visualization, the function *RunUMAP* (20 dimensions) was performed. For GSE128423, 6 files corresponding to stromal cells (GSM3674224, GSM3674225, GSM3674226, GSM3674227, GSM3674228, GSM3674229) were downloaded and read into Seurat (versions 4 and 5) in R (version 4.3.1) then merged into a single object. Low quality cells were filtered according to the following criteria: mitochondrial reads per cell >5.5%, number of reads per cell >65,000, number of unique genes per cell <200 and >7000. The merged object was then processed using the following functions: *NormalizeData*, *FindVariableFeatures*, *ScaleData*, *RunPCA* (30 dimensions), *RunUMAP* (20 dimensions), *FindNeighbors* (20 dimensions), *FindClusters* (resolution 0.5). Since the original cell annotation was not available in the submission, the top 50 differentially expressed genes for each cluster were obtained from *Supplementary Data 1* of Baryawno et al. (2019), and the function *SCINA* from the R package SCINA (Zhang et al., 2019) was used to predict the identity of every cell, using the same nomenclature shown in *Figure 1* of Baryawno et al. (2019). For GSE145477, the count tables derived from 1 month old (GSM4318799) and 1.5 month old (GSM4318800) mice were downloaded and used to create a merged Seurat object. The cells were then filtered and processed as described for GSE128423. For creation of the 10X mesenchymal atlas, clusters corresponding to endothelial cells or hematopoietic cells were manually identified in each component dataset using characteristic marker genes based on the *FindAllMarkers* functions, and these cells were removed. The raw counts for the remaining mesenchymal cells were normalized separately using *SCTransform* then integrated using SCT integration functions (*SelectIntegrationFeatures*, *PrepSCTIntegration*, *FindIntegrationAnchors*, *IntegrateData*). The integrated assay was then processed using *RunPCA* (30 components), *RunUMAP* (20 dimensions), *FindNeighbors* (20 dimensions), and *FindClusters* (resolution 0.5). For trajectory inference, functions from Monocle3 were implemented through SeuratWrappers (Trapnell et al., 2014). The *learn_graph* and *order_cells* functions were run twice using two different starting nodes corresponding to different MSC populations; LepR^+^ MSCs and Sca-1^+^ MSCs points were defined using genelists from previous publications with the *AddModuleScore* function. Other 10X and Smart-seq datasets were mapped to the 10X mesenchymal atlas using *FindTransferAnchors* and *MapQuery* functions. To align stromal nomenclature, datasets from Wolock et al., 2019 [GSE132151] and Tikhonova et al., 2019 [GSE108892] were also analyzed in this study. For GSE132151, counts and metadata were downloaded and imported into Seurat, and processed as for GSE128423, except that cell annotations from the original study were already available in the metadata. For GSE108892, the count tables and metadata for 5 samples corresponding to steady state and the controls for the 5FU experiments were downloaded (GSM2915575, GSM2915576, GSM2915577, GSM2915578, GSM2915579) and merged into a single Seurat object. This object was processed as for GSE128423, except that the original annotation of cells was available in the author metadata. All these datasets were used for mapping between other datasets using the *FindTransferAnchors* and *MapQuery* functions.

### Spearman correlations

The *clustify* function in the R package ClustifyR was used to generate spearman correlations for gene expression between Smart-seq stromal populations or 10X mesenchymal subclusters and published bulk RNA sequencing or microarray data, using the top 2,000 highly variable genes from the scRNA-seq datasets. For bulk RNA sequencing data from Asada et al., 2017 [GSE89811], Xu et al., 2018 [GSE104701], Helbling et al., 2019 [GSE133922], He et al., 2017 [GSE98587], and Tikhonova et al., 2019 [GSE108892], tables of normalized gene counts were downloaded directly and used as input for spearman correlations. For Mamidi et al., 2023 [GSE143249], Fastq files were downloaded, trimmed with TrimGalore, and aligned to the murine mm10 genome using Salmon to generate a normalized counts table. For microarray data contained in Greenbaum et al., 2013 [GSE43613], Worthley et al., 2015 [GSE57729], Yu et al., 2015 [GSE66042], and Ding et al., 2012 [GSE33158], raw data were downloaded and normalized using the *rma* function in the R package oligo (Carvalho & Irizarry, 2010). Gene annotations were added using the *annotateEset* function (affycoretools package; MacDonald, 2024) and the appropriate reference for the microarray, before rows corresponding to the same genes were collapsed with the *collapseRows* function (WGCNA package, Langfelder & Horvath, 2008). Output from clustify was scaled between 0 and 1 across stromal populations or mesenchymal subclusters.

### Stromal colony assays

For CFU-F assays, Endo MSC-S cells (15-300 cells) and mMPr/MPr (170-300 cells) were sorted directly into 6-well plates containing 1.5 ml αMEM supplemented with 10% FBS, 100 U/ml penicillin/100 μg/ml streptomycin, and 50 μM 2-mercaptoethanol. Cells were cultured under normal oxygen level for 11 days before staining with Giemsa-Wright to score colonies of 25 or more cells. CM MSC-L cells (100-300 cells) or (i)MSC-L (1,000 cells) were sorted directly into 6-well plates containing 1.5 mL of DMEM supplemented with 20% FBS, 100 U/ml penicillin/100 μg/ml streptomycin, 50 μM 2-mercaptoethanol, and 10 μM ROCK inhibitor (iROCK, Tocris, Y-27632). Cells were cultured in hypoxic conditions (5% O_2_) for 8 days before staining with Giemsa-Wright to score colonies of 25 or more cells. For osteoblastic differentiation, αMEM or DMEM based medium were further supplemented with 3 mM β-glycerol phosphate (Sigma G9891) and 50 μg/mL ascorbic acid-2-phosphate (Sigma A8950). To stain for alkaline phosphatase, MSC-S cells (15-300 cells), mMPr/MPr (170-300 cells), or MSC-L cells (100-300 cells) were sorted directly into 6-well plates containing 1.5 ml of their respective culture medium, which was replaced with osteoblastic differentiation medium 2 days later. The cells were cultured for a further 9 days (until day 11 after isolation) and fixed with 1:10 phosphate-buffered formalin (Fisher scientific, 23-245-684) for 10 minutes. The cells were stained for 1 hour with 0.1 M Tris-HCl, pH 8.3 containing 0.6 mg/ml Fast Red Violet LB Salt (Sigma, F3381), 0.1 mg/ml Naphthol AS-MX phosphate (Sigma, N4875), and 0.4% N,N-dimethylformamide before counting CFU-ALP colonies of at least 25 cells. For Von Kossa staining, the cell culture was maintained in osteoblastic differentiation medium until day 23, fixed in phosphate-buffered formalin for 10 minutes, and then stained with 1% silver nitrate solution (Electron Microscopy Sciences 26212-01) for 20 minutes with light exposure, and rinsed with sodium thiosulfate (Ricca Chemical R7866500). For adipocytic differentiation and Oil-Red-O staining, MSC-S cells (15-300 cells), mMPr/MPr (170-300 cells), and MSC-L cells (100-300 cells) were sorted directly into 6-well plates containing 1.5 ml culture medium and cultured in the appropriate conditions until day 11 or day 8, respectively. The medium was then replaced with StemPro Adipogenesis Differentiation Kit (ThermoFisher). Four days later, the cells were fixed in phosphate-buffered formalin for 1 hour or longer and stained with Oil-Red-O (O1391) for 10 minutes before counting CFU-ORO colonies of at least 25 cells.

### Complete blood cell counts

Blood was collected by cardiac puncture after euthanasia and immediately transferred into EDTA-coated tubes (Greiner Bio-One, 0.5 ml). Complete blood cell counts were performed with an Oxford Science Genesis analyzer.

### BM section imaging

Whole femurs from mice injected with MLL/AF9 blasts or control mice were dissected, placed in plastic mounts containing OCT, and immediately frozen at -80°C. Sections of 7 μm thickness were cut using a Leica CM3050S cryostat with tungsten carbide blade and transferred to glass slides using a CryoJane tape transfer system (Leica). Slides were dried for 20 minutes at room temperature then fixed in acetone at -20°C for 10 minutes. When dry, a circle was drawn around tissue sections using a PAP pen (Ted Pella) before washing three times for 5 minutes with PBS. Slides were blocked with 10% goat serum (ThermoFisher Scientific) in PBS for 1 hour at room temperature and then stained with rabbit anti-mouse laminin primary antibody (Sigma-Aldrich, 1:100 dilution in PBS with 10% goat serum) for 2 hours at room temperature. After this, slides were washed three times for 5 minutes with PBS, then stained with goat anti-rabbit IgG AF647 secondary antibody (Invitrogen, 1:1000 dilution) and DAPI (5 μg/ml) in PBS with 10% goat serum for 90 minutes at room temperature. Slides were washed as before then mounted with ProLong Glass Antifade Mountant (ThermoFisher Scientific) and a coverslip was applied. Images were acquired with a SP8 inverted confocal microscope (Leica) with a 20X objective lens using Leica Application Suite X. A z-stack of 1 μm was acquired across sections, and final images are maximum intensity projections.

### BM fluid isolation and assays

To isolate BM fluid, 2 tibiae from each mouse were flushed 15 times each with the same 350 μl of HBSS with 2% FBS into 1.5 ml tubes. Tubes were then centrifuged at 1,000 *g* for 10 minutes, and the supernatant was removed into new tubes and frozen at -80°C. Concentrations of chemokines were measured in BM fluid using the LEGENDplex mouse proinflammatory chemokine panel (13plex, BioLegend) with a V bottom plate according to the kit manual. Standards were diluted in matrix B, and samples were diluted 1:2 in assay buffer. Data were acquired with a Novocyte Penteon (Agilent).

### Transwell assays

To evaluate chemoattractant activity of BM fluid from PBS or 3 day-pIC-exposed mice, 50 μl of BM fluid was diluted with 750 μl of complete IMDM medium (containing 5% FBS, 100 U/ml penicillin and 100 μg/ml streptomycin, 1X non-essential amino acids, 1 mM sodium pyruvate, 1X GlutaMAX, and 50 μM 2-mercaptoethanol), and placed in the bottom of a 24 well plate. A transwell insert (6.5 mm diameter, 5 μm pores) was then placed into the well. Whole BM was obtained from *Ifnar-/-* mice by flushing two femurs with a 3 ml syringe and a 21G needle and performing ACK RBC lysis. 10^6^ whole BM cells were placed into the top of the transwell insert in a volume of 100 μl of complete IMDM medium, and the plate was incubated at 37°C with 5% CO_2_ for 2 hours. After this, the transwell insert was removed and the medium in the well was re-suspended and transferred to 1.5 ml tubes, centrifuged, and stained with an antibody mixture of Ly6G-FITC (BioLegend, 127606, 1:400), Ly6C-APC/Cy7 (BioLegend, 128026, 1:400), CD8-AF700 (BD, 557959, 1:400), B220-BV421 (BioLegend, 103240, 1:800), CD4-BV510 (BioLegend, 100553, 1:400), CD11b-PE/Cy7 (Invitrogen, 25-0112-82, 1:800), CD115-PE (BioLegend, 135506, 1:100), CD3-APC (BioLegend, 100220, 1:100), and NK1.1-PE/Cy5 (BioLegend, 108716, 1:400) for 30 minutes on ice. Cells were then washed and re-suspended in HBSS with 2% FBS with 1 μg/ml propidium iodide before acquisition on a Novocyte Quanteon or Penteon (Agilent).

### qRT-PCR analyses

Approximately 10,000 cells per population were sorted directly by FACS into RLT plus lysis buffer (Qiagen) with 2-mercaptoethanol and stored at -80 °C until purification with a RNeasy Plus Micro Kit (Qiagen) according to the manufacturer’s protocol. Following column purification, RNA was immediately reverse transcribed using a SuperScriptIII kit with random hexamers (ThermoFisher Scientific). qRT-PCR runs were performed on a QuantStudio 7 Flex Real-Time PCR system (Applied Biosystems) using SYBR Green reagents (Bio-Rad), the cDNA equivalent of 200 cells per reaction, and triplicate technical measurements per biological repeat. Cycle threshold values were normalized to *Actb*.

### Statistics and reproducibility

All experiments were repeated as indicated; n indicates the numbers of independent biological repeats. Data are expressed as the mean ± s.d. unless otherwise indicated. Mice used for treatment were assigned to experimental groups based on age and genotype, randomized with respect to sex, and samples were alternated whenever possible. Data collection and analysis were not performed blind to the conditions of the experiments. Data were processed using Microsoft Excel (v.16), and statistical significance was evaluated using GraphPad Prism (v.10). Figures were made using GraphPad Prism. Data distribution was assumed to be normal, but this was not formally tested.

### Data and Software Availability

Data sets that support the findings of this study have been deposited in the Gene Expression Omnibus (GSE276784). Source data for all the figures are provided with the paper. All other data are available from the corresponding author upon reasonable request.

