## Extended Data Figures 1 to 10 and Table 1 to 3 for "Inflammation perturbs hematopoiesis by remodeling specific compartments of the bone marrow niche"

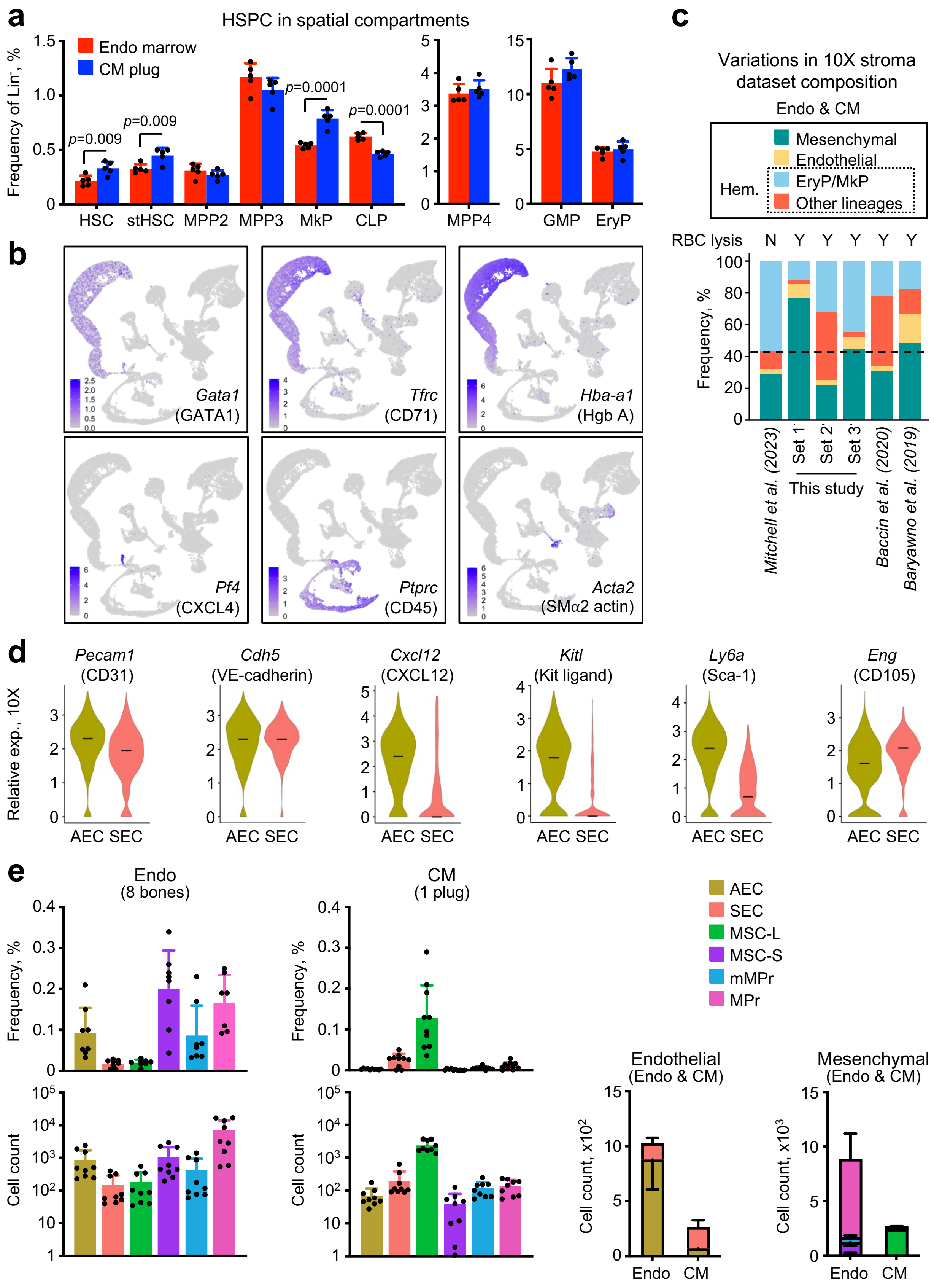
a
HSPC in spatial compartments
Endo marrow
CM plug
Frequency of Lin⁻, %
p=0.009
p=0.009
p=0.0001
p=0.0001
HSC
stHSC
MPP2
MPP3
MkP
CLP
MPP4
GMP
EryP
b
Gata1 (GATA1)
Tfrc (CD71)
Hba-a1 (Hgb A)
Pf4 (CXCL4)
Ptprc (CD45)
Acta2 (SMα2 actin)
c
Variations in 10X stroma dataset composition
Endo & CM
Mesenchymal
Endothelial
EryP/MkP
Other lineages
Hem.
RBC lysis N Y Y Y Y Y
Frequency, %
Mitchell et al. (2023)
Set 1
Set 2
Set 3
This study
Baccin et al. (2020)
Baryawno et al. (2019)
d
Pecam1 (CD31)
Cdh5 (VE-cadherin)
Cxcl12 (CXCL12)
Kitl (Kit ligand)
Ly6a (Sca-1)
Eng (CD105)
Relative exp., 10X
AEC SEC
e
Endo (8 bones)
CM (1 plug)
Frequency, %
Cell count
AEC
SEC
MSC-L
MSC-S
mMPr
MPr
Endothelial (Endo & CM)
Mesenchymal (Endo & CM)
Cell count, x10²
Cell count, x10³
Endo
CM

**Extended Data Figure 1 | Spatial compartmentation of stromal and hematopoietic cells. a**, Frequency of indicated hematopoietic stem and progenitor cells (HSPC) among lineage negative BM cells in central marrow (CM) and endosteal (Endo) fractions. HSC: hematopoietic stem cell; stHSC: short-term HSC; MPP2/3/4: multipotent progenitor 1/2/3; MkP: megakaryocyte progenitor; CLP: common lymphoid progenitor; GMP: granulocyte macrophage progenitor; EryP: erythroid progenitor. **b**, Feature plots showing expression of indicated genes in the 10X stroma map. **c**, Frequency of indicated cell types in stroma scRNA-seq datasets from this study (Set 1-3), and from previously published datasets with or without red blood cell (RBC) lysis. Hem.: hematopoietic cells, Y: yes, N: no. **d**, Expression of surface marker genes associated with arterial endothelial cell (AEC) and sinusoidal endothelial cell (SEC) cluster identity in the 10X stroma map. **e,** Frequency (top) and absolute number (bottom) of major stromal cell populations in CM and Endo preparations. MSC-L: $LepR^+$ mesenchymal stromal cell, MSC-S: Sca-$1^+$ mesenchymal stromal cell, (m)MPr: (multipotent) mesenchymal progenitor. Data in (a) and (e) are means ± S.D. with points showing values for individual mice; *P. values*, Student's t test. Data in (d) are violin plots of relative expression (exp.) of 10X SCT transformed counts with median.

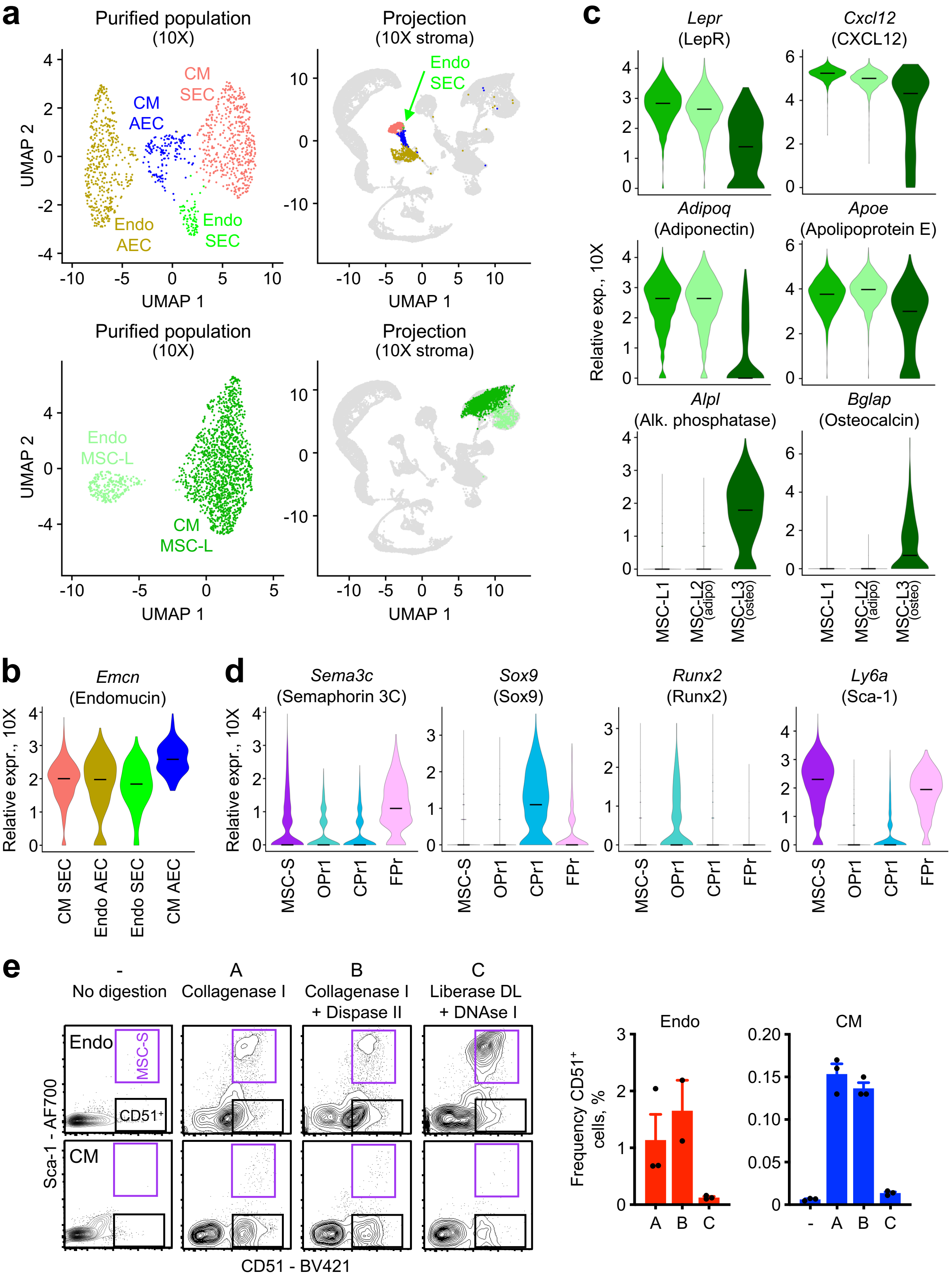
a
Purified population (10X)
CM SEC
CM AEC
Endo AEC
Endo SEC
UMAP 2
UMAP 1
Projection (10X stroma)
Endo SEC
Endo MSC-L
CM MSC-L
b
Emcn (Endomucin)
Relative expr., 10X
CM SEC
Endo AEC
Endo SEC
CM AEC
c
Lepr (LepR)
Cxcl12 (CXCL12)
Adipoq (Adiponectin)
Apoe (Apolipoprotein E)
Alpl (Alk. phosphatase)
Bglap (Osteocalcin)
Relative exp., 10X
MSC-L1
MSC-L2 (adipo)
MSC-L3 (osteo)
d
Sema3c (Semaphorin 3C)
Sox9 (Sox9)
Runx2 (Runx2)
Ly6a (Sca-1)
MSC-S
OPr1
CPr1
FPr
e
- No digestion
A Collagenase I
B Collagenase I + Dispase II
C Liberase DL + DNAse I
Endo
CM
MSC-S
CD51⁺
Sca-1 - AF700
CD51 - BV421
Frequency CD51⁺ cells, %

**Extended Data Figure 2 | Optimization of stromal isolation approach. a**, UMAP dimension reduction (left) and projection onto the reference 10X stroma map (right) of purified SEC/AEC (top) and MSC-L (bottom) that were oligo-hashed and isolated from both CM and Endo compartments from young mice (n = 1 biological replicate, 7 male mice) and analyzed by 10X scRNA-seq. **b**, Differential expression of *Emcn* (endomucin) gene in purified CM and Endo SEC/AEC. **c**, Expression of signature genes associated with MSC-L cluster identity in 10X stroma map. Adipo: adipocyte bias, Osteo: osteoblast biased. **d**, Expression of signature genes associated with MSC-S and mesenchymal progenitor cluster identity in the 10X stroma map. OPr: osteoprogenitor, CPr: chrondroblast progenitor, FPr: fibroblast progenitor. **e**, Representative flow cytometry plots (left) and quantification of $CD51^{+}$ cells isolated from CM and Endo preparations using no digestion or the indicated digestion protocols (A-C). Data in (b), (c), and (d) are violin plots of relative expression (exp.) of 10X SCT transformed counts with median. Data in (e) are means ± S.D. with points showing values for individual mice.

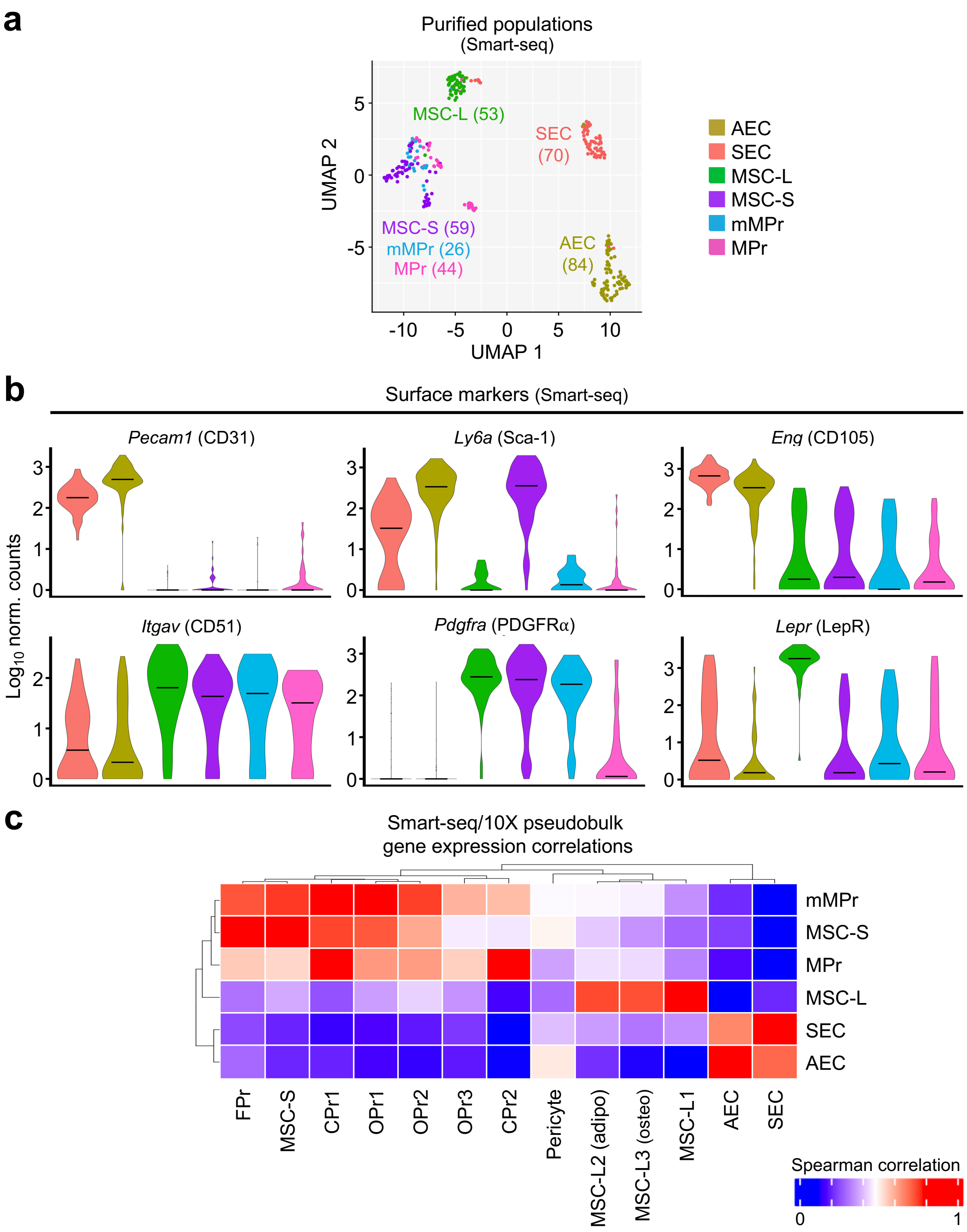
a
Purified populations
(Smart-seq)
UMAP 1
UMAP 2
MSC-L (53)
SEC
(70)
MSC-S (59)
mMPr (26)
MPr (44)
AEC
(84)
AEC
SEC
MSC-L
MSC-S
mMPr
MPr
b
Surface markers (Smart-seq)
Pecam1 (CD31)
Ly6a (Sca-1)
Eng (CD105)
Itgav (CD51)
Pdgfra (PDGFRα)
Lepr (LepR)
Log10 norm. counts
c
Smart-seq/10X pseudobulk
gene expression correlations
mMPr
MSC-S
MPr
MSC-L
SEC
AEC
FPr
MSC-S
CPr1
OPr1
OPr2
OPr3
CPr2
Pericyte
MSC-L2 (adipo)
MSC-L3 (osteo)
MSC-L1
AEC
SEC
Spearman correlation
0
1

**Extended Data Figure 3 | Molecular characterization of major stromal populations. a**, UMAP dimension reduction of purified major CM (SEC, MSC-L) and Endo (MSC-S, mMPr, MPr) stromal cells analyzed by Smart-seq and previously published in Mitchell et al. (2023). **b,** Expression of surface marker genes used to isolate major CM and Endo stromal cells for Smart-seq analyses. Data are violin plots of Log10 normalized (norm.) Smart-seq counts with median. **c,** Heatmap showing row-normalized Spearman correlations between indicated stromal populations isolated for Smart-seq analyses with pseudobulk profiles from clusters identified in the 10X stroma map.

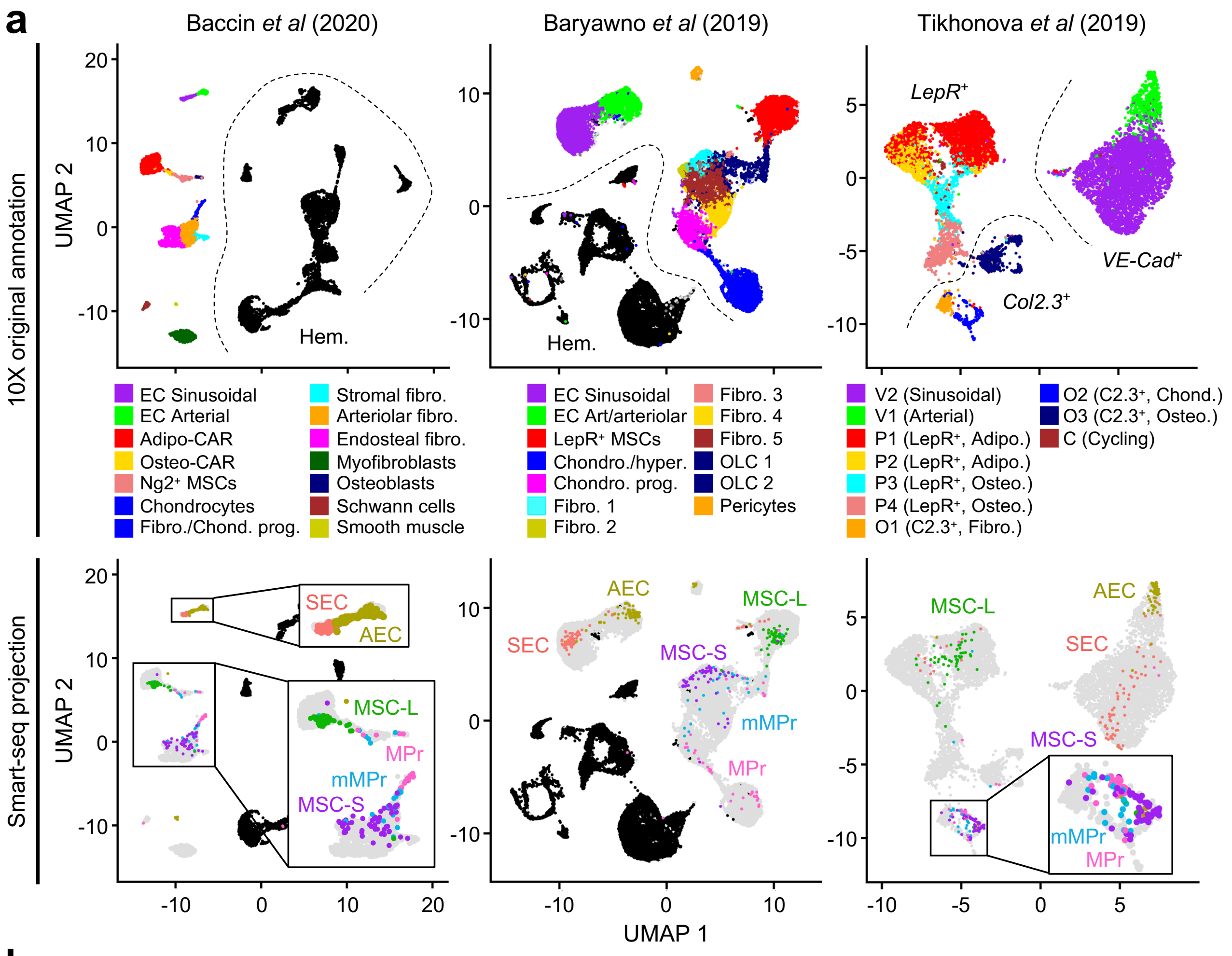


**b**

| This study (Flow) | This study (10X) | Baccin *et al* (2020) | Baryawno *et al* (2019) | Tikhonova *et al* (2019) |
|---|---|---|---|---|
| MSC-L | MSC-L1<br>MSC-L2 (adipo)<br>MSC-L3 (osteo) | Adipo-CAR<br>Osteo-CAR | LepR+ MSC | P1 (LepR+, Adipo.)<br>P2 (LepR+, Adipo.) |
| MSC-S | MSC-S<br>FPr | Endosteal fibroblast<br>Arteriolar fibroblast<br>Stromal fibroblast | Fibroblast 1<br>Fibroblast 2<br>Fibroblast 3 | O2 (Col2.3+, Chondro.) |
| mMPr | OPr1<br>CPr1 | Fibro./Chondro. progen. | OLC 1<br>Fibroblast 5<br>Chondroblast progen. | O2 (Col2.3+, Chondro.) |
| MPr | OPr2<br>OPr3<br>CPr2 | Chondrocytes<br>Osteoblasts | Chondroblasts<br>Chondroblast progen.<br>OLC 1<br>OLC 2 | O2 (Col2.3+, Chondro.) |
| SEC | SEC | EC Sinusoidal | EC Sinusoidal | V2 (Sinusoidal) |
| AEC | AEC | EC Arterial | EC Arterial/Arteriolar | V1 (Arterial) |

**Extended Data Figure 4 | Alignment of stromal nomenclature across studies. a**, Projection of Smart-seq stromal cell transcriptomes onto the indicated previously published 10X stromal datasets, with original annotations derived from those studies. Hematopoietic cell (Hem.) contamination is shown in black and the fluorescence reporter mice lines used to identify stroma element is provided for Tikhonova et al. (2019). Of note, Baccin et al. (2020) used undigested stromal preparation likely preserving isolation of Schwann and smooth muscle cells. **b**, Alignment of names for major stromal populations between this study and previous publications.

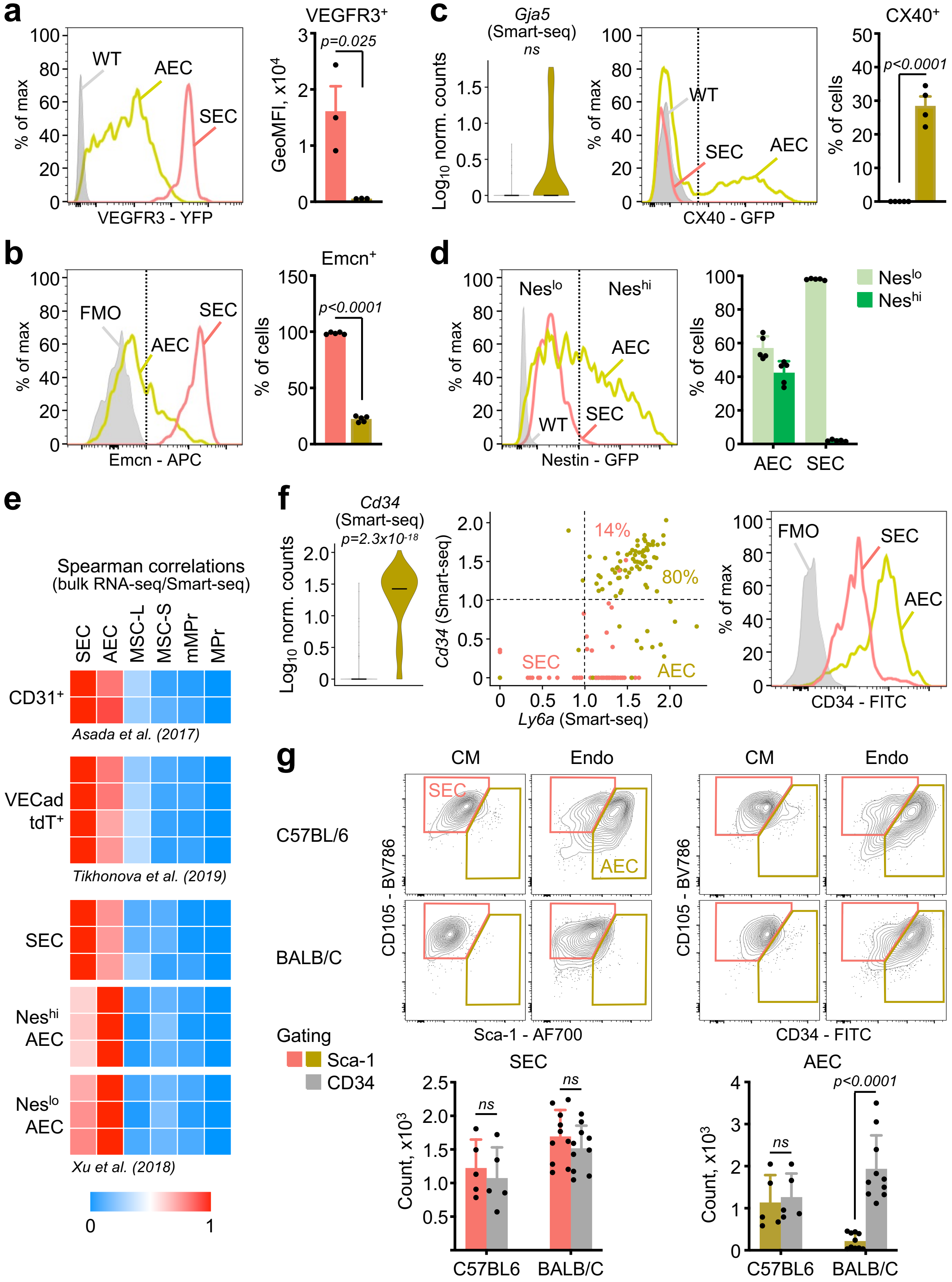

a
VEGFR3 - YFP
% of max
WT
AEC
SEC
VEGFR3⁺
GeoMFI, x10⁴
p=0.025
b
Emcn - APC
FMO
Emcn⁺
% of cells
p<0.0001
c
Gja5 (Smart-seq)
ns
Log10 norm. counts
CX40 - GFP
CX40⁺
d
Nestin - GFP
Neslo
Neshi
e
Spearman correlations (bulk RNA-seq/Smart-seq)
SEC
AEC
MSC-L
MSC-S
mMPr
MPr
CD31⁺
Asada et al. (2017)
VECad tdT⁺
Tikhonova et al. (2019)
Neshi AEC
Neslo AEC
Xu et al. (2018)
f
Cd34 (Smart-seq)
p=2.3x10-18
Cd34 (Smart-seq)
Ly6a (Smart-seq)
14%
80%
CD34 - FITC
g
CM
Endo
C57BL/6
BALB/C
CD105 - BV786
Sca-1 - AF700
Gating
Sca-1
CD34
Count, x10³
C57BL6

**Extended Data Figure 5 | Functional comparison of CM SECs and Endo AECs. a**, Representative flow cytometry plots (left) and quantification of geometric mean fluorescence intensity (GeoMFI, right) of VEGFR3 expression in SECs and AECs of *Vegfr3-Yfp* reporter mice. **b**, Representative flow cytometry plots (left) and quantification (right) of the proportion of SECs and AECs stained with Endomucin antibody ($Emcn^+$ cells). **c**, Expression of *Gja5* in Smart-seq SECs and AECs (left) with representative flow cytometry plots (middle) and quantification (right) of the proportion of SECs and AECs expressing connexin 40 (CX40) in *Cx40-Gfp* reporter mice. **d**, Representative flow cytometry plots (left) and quantification (right) of the proportion of SECs and AECs with low ($Nes^{lo}$) or high ($Nes^{hi}$) Nestin expression in *Nestin-Gfp* reporter mice. **e**, Spearman correlation between Smart-seq stromal populations with bulk RNA sequencing data for the indicated endothelial cell types obtained from previous publications. VECad tdT: VE-cadherin tdTomato. **f**, Expression of CD34 (*Cd34*) in Smart-seq SECs and AECs alone (left) or compared to Sca-1 (*Ly6a*) (middle) with representative flow cytometry plots of CD34 antibody staining in SECs and AECs. **g**, Representative flow cytometry plots showing identification (top) and quantification (bottom) of the number of SECs and AECs identified with either Sca-1 (left) or CD34 (right) antibody staining in C57BL/6 and BALB/C mice. WT: wild type staining control included for the analyses of fluorescent reporter mice in (a), (c), and (d), FMO: fluorescence minus one control included for the analyses of antibody staining in (b) and (f). Data in (c, left) and (f, left) are violin plots of Log10 normalized (norm.) Smart-seq counts with median. Other data except for (e) and (f) are means ± S.D. with points showing values for individual mice; *P. values* for (a), (b), and (c), Student's t test; *P. values* for (d) and (g), two-way ANOVA with Sidak's *post hoc* test; *P. values* for (c) and (f), Wilcoxon rank sum test.

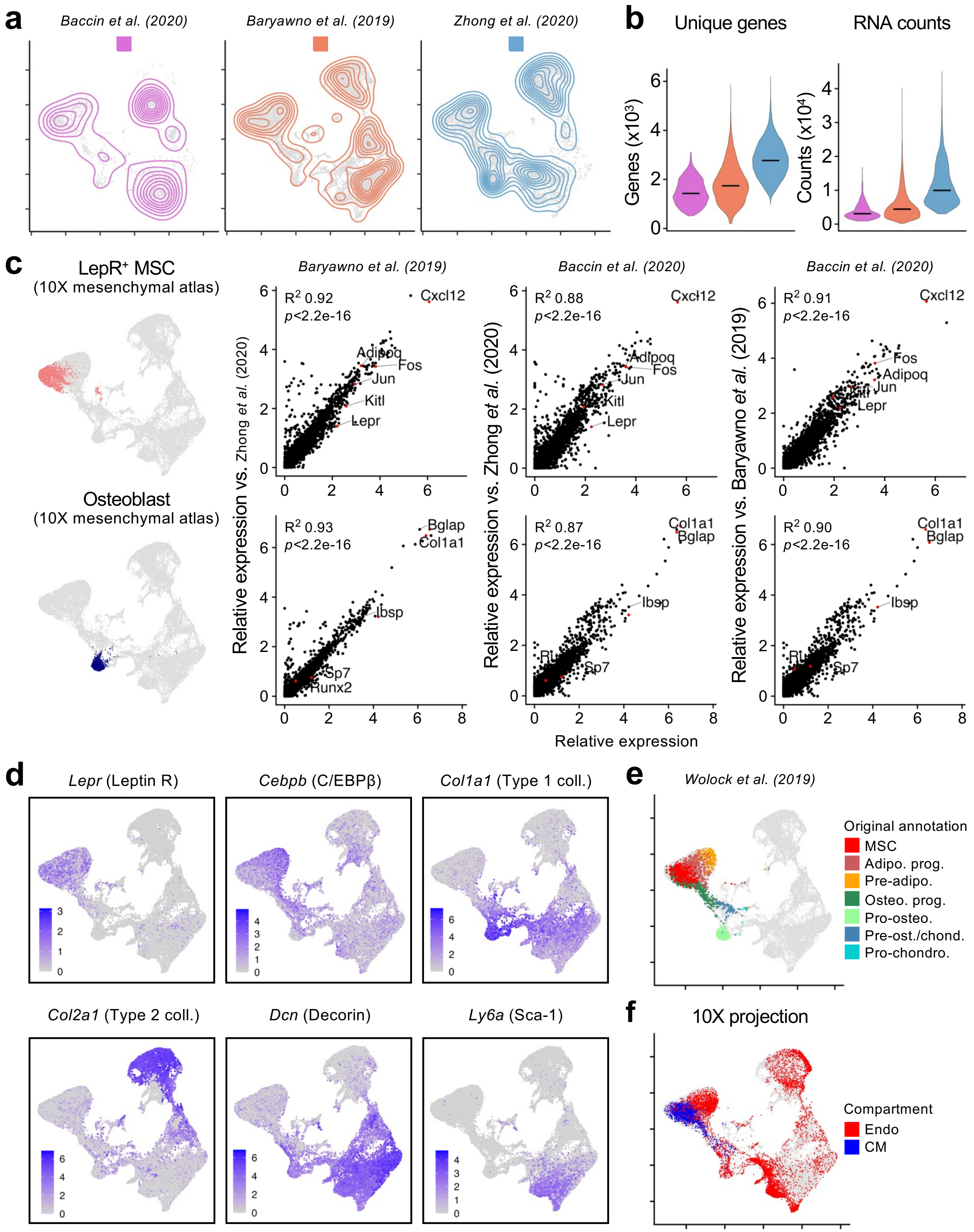
a
Baccin et al. (2020)
Baryawno et al. (2019)
Zhong et al. (2020)
b
Unique genes
RNA counts
Genes (x10³)
Counts (x10⁴)
c
LepR⁺ MSC
(10X mesenchymal atlas)
Osteoblast
(10X mesenchymal atlas)
Baryawno et al. (2019)
Baccin et al. (2020)
Baccin et al. (2020)
R² 0.92
p<2.2e-16
R² 0.88
p<2.2e-16
R² 0.91
p<2.2e-16
R² 0.93
p<2.2e-16
R² 0.87
p<2.2e-16
R² 0.90
p<2.2e-16
Cxcl12
Adipoq
Fos
Jun
Kitl
Lepr
Bglap
Col1a1
Ibsp
Sp7
Runx2
Relative expression vs. Zhong et al. (2020)
Relative expression vs. Zhong et al. (2020)
Relative expression vs. Baryawno et al. (2019)
Relative expression
d
Lepr (Leptin R)
Cebpb (C/EBPβ)
Col1a1 (Type 1 coll.)
Col2a1 (Type 2 coll.)
Dcn (Decorin)
Ly6a (Sca-1)
e
Wolock et al. (2019)
Original annotation
MSC
Adipo. prog.
Pre-adipo.
Osteo. prog.
Pro-osteo.
Pre-ost./chond.
Pro-chondro.
f
10X projection
Compartment
Endo
CM

**Extended Data Figure 6 | Construction of the 10X mesenchymal atlas. a**, UMAPs of individual 10X scRNA-seq datasets derived from the indicated previous publications that were integrated to form the composite 10X mesenchymal atlas. **b**, Number of unique genes and gene counts from color-coded individual datasets, with medians. **c**, Correlations across individual datasets in gene expression for 2 examples of 10X mesenchymal atlas clusters, with selected gene names (red dots) and linear regression $R^2$ values. **d**, Feature plots showing expression of indicated genes in the 10X mesenchymal atlas. **e**, Projection of previously annotated 10X stromal cell transcriptomes onto the 10X mesenchymal atlas, with original annotation: adipo.: adipocyte, osteo.: osteoblast; chond.: chondroblast. **f**, Projection of 10X stroma transcriptomes from this study onto the 10X mesenchymal atlas, colored by spatial compartment as CM (blue) and Endo (red).

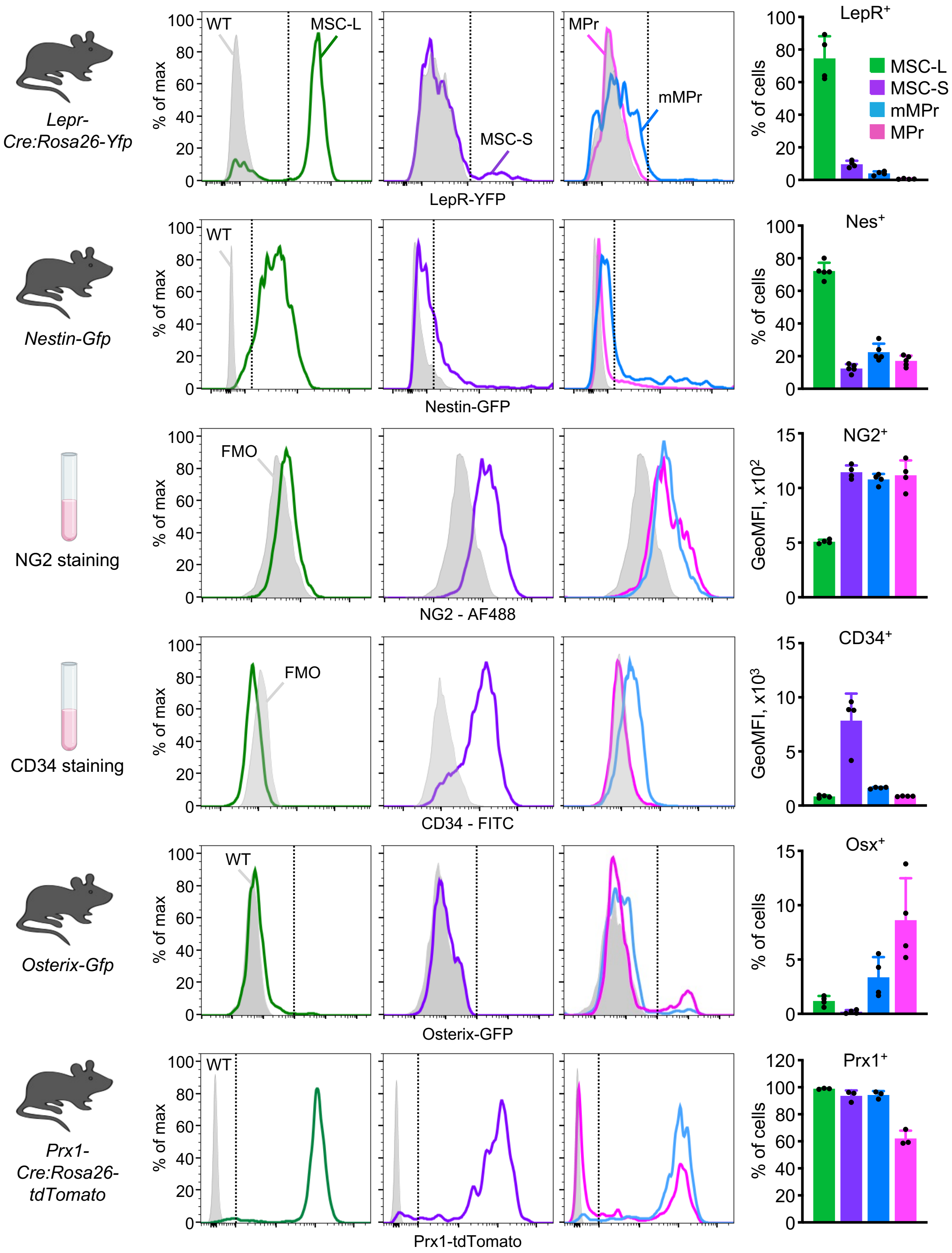

Lepr-Cre:Rosa26-Yfp
Nestin-Gfp
NG2 staining
CD34 staining
Osterix-Gfp
Prx1-Cre:Rosa26-tdTomato
% of max
WT
MSC-L
MSC-S
MPr
mMPr
FMO
LepR-YFP
Nestin-GFP
NG2 - AF488
CD34 - FITC
Osterix-GFP
Prx1-tdTomato
LepR+
Nes+
NG2+
CD34+
Osx+
Prx1+
% of cells
GeoMFI, x10²
GeoMFI, x10³

**Extended Data Figure 7 | Expression of fluorescent and protein markers among major stromal cell types.** Schematic of the used fluorescent reporter mice or antibody staining (left), representative flow cytometry plots (middle), and quantification of expression in CM MSC-L and Endo MSC-S, mMPr and MPr. Wild type (WT) staining controls are included for the analyses of reporter mice and results are expressed as proportion of cells expressing the fluorescent marker. Fluorescence minus one (FMO) controls are included for the analyses of antibody staining and results are expressed as geometric mean fluorescence intensity (GeoMFI) measurements. Nes: nestin, Osx: osterix. Data are means ± S.D. with points showing values for individual mice.

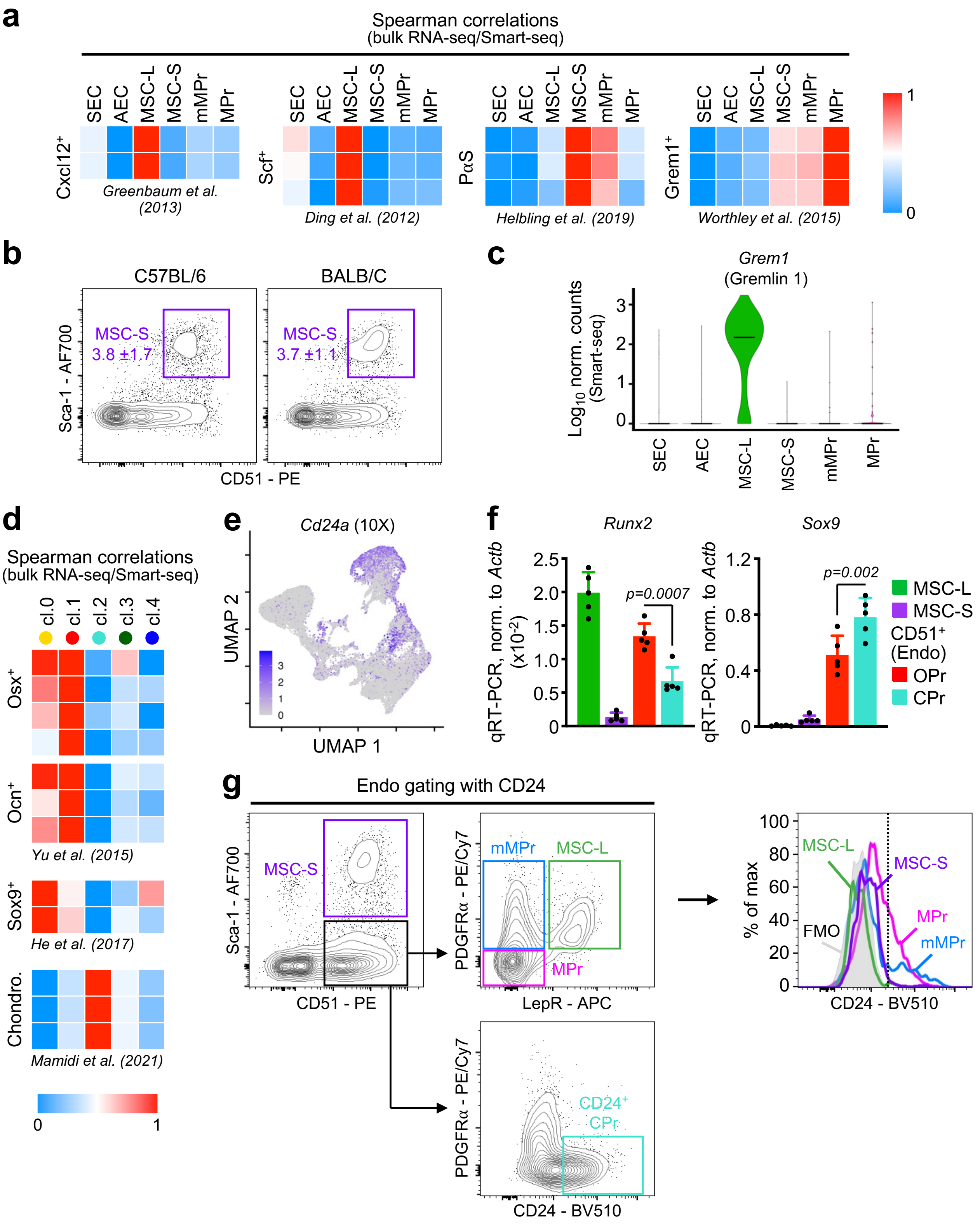
a
Spearman correlations
(bulk RNA-seq/Smart-seq)
SEC AEC MSC-L MSC-S mMPr MPr
Cxcl12+
Greenbaum et al. (2013)
Scf+
Ding et al. (2012)
PαS
Helbling et al. (2019)
Grem1+
Worthley et al. (2015)
1
0
b
C57BL/6
BALB/C
MSC-S 3.8 ±1.7
MSC-S 3.7 ±1.1
Sca-1 - AF700
CD51 - PE
c
Grem1
(Gremlin 1)
Log10 norm. counts (Smart-seq)
d
Spearman correlations
(bulk RNA-seq/Smart-seq)
cl.0 cl.1 cl.2 cl.3 cl.4
Osx+
Ocn+
Yu et al. (2015)
Sox9+
He et al. (2017)
Chondro.
Mamidi et al. (2021)
e
Cd24a (10X)
UMAP 2
UMAP 1
f
Runx2
Sox9
qRT-PCR, norm. to Actb (x10-2)
qRT-PCR, norm. to Actb
p=0.0007
p=0.002
MSC-L
MSC-S
CD51+ (Endo)
OPr
CPr
g
Endo gating with CD24
MSC-S
Sca-1 - AF700
CD51 - PE
mMPr
MSC-L
MPr
PDGFRα - PE/Cy7
LepR - APC
CD24+ CPr
CD24 - BV510
% of max
FMO
CD24 - BV510

**Extended Data Figure 8 | Refined characterization of Endo mesenchymal progenitors. a**, Spearman correlation between Smart-seq stromal populations with bulk RNA sequencing data for the indicated cell types obtained from previous publications. PαS: PDGFR-$\alpha^+$/Sca-$1^+$ cells. Grem1: Gremlin 1. **b**, Representative flow cytometry plots of Sca-1 staining in Endo $CD51^+$ fraction in C57BL/6 and BALB/C mice, with quantification of MSC-S frequency (n = 5 individual mice). **c**, Differential expression of *Grem1* in purified Smart-seq populations. Results are shown as violin plots of Log10 normalized (norm.) Smart-seq counts with median. **d**, Spearman correlation between identified Smart-seq PHATE Louvain clusters with bulk RNA sequencing data for the indicated cell types obtained from previous publications. Osx: osterix, Ocn: osteocalcin, Chondro.: primary chondrocytes. **e**, Feature plots showing expression of *Cd24a* in the 10X mesenchymal atlas. **f**, *Runx2* and *Sox9* expression measured by qRT-PCR in the indicated stroma populations isolated by flow cytometry. OPr: $CD24^-$ subset of the $CD51^+$ Endo fraction, CPr: $CD24^+$ subset of the $CD51^+$ Endo fraction. Results are normalized to expression of *Actb* and data are means ± S.D. with points showing values for individual mice; *P. values*, one-way ANOVA with Tukey *post-hoc* test. **g**, Representative flow cytometry plots showing Endo gating with CD24 for CPr (left) and expression of CD24 in major Endo stromal populations (right).

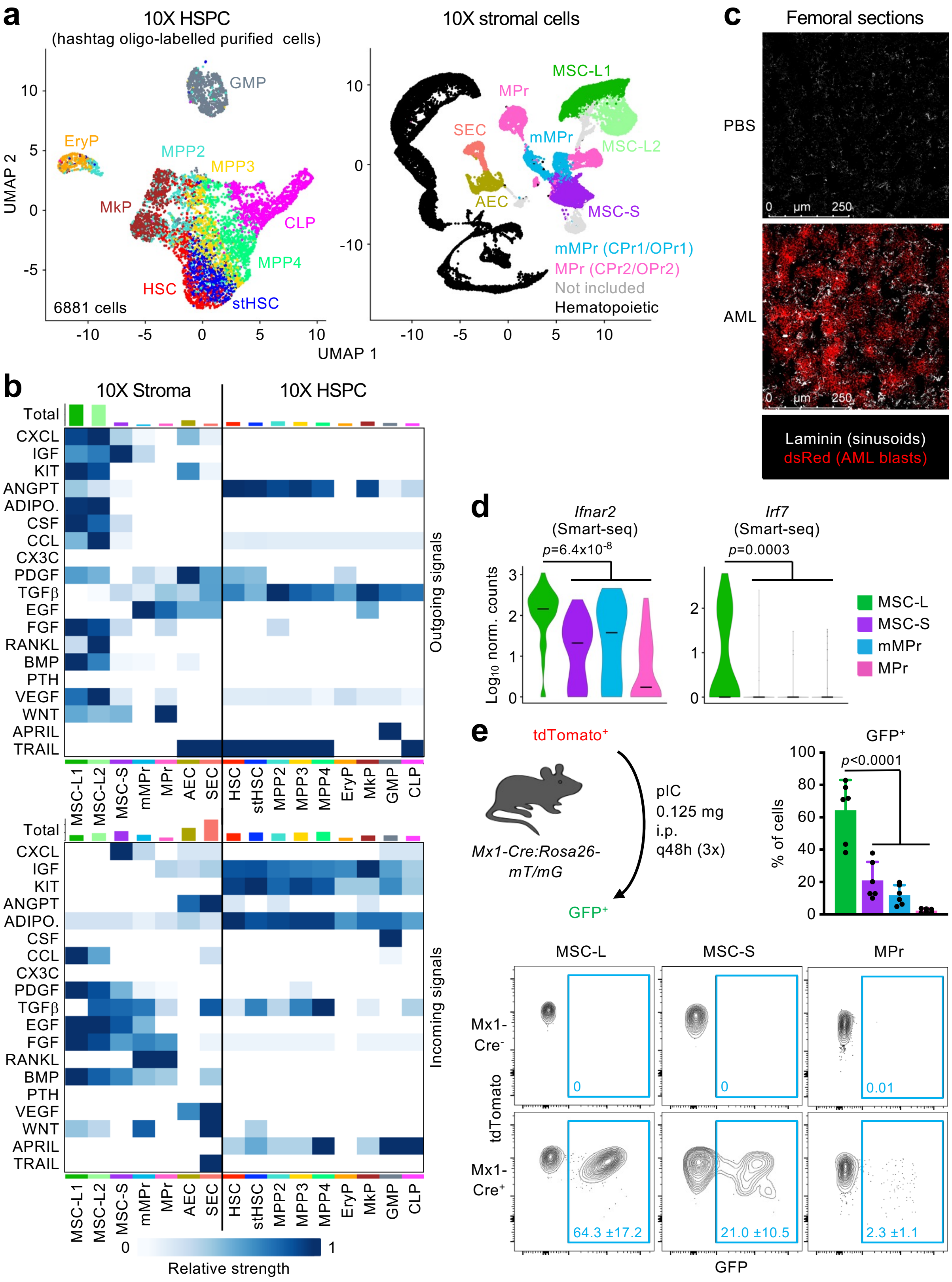
a
10X HSPC
(hashtag oligo-labelled purified cells)
GMP
EryP
MPP2
MPP3
MkP
CLP
MPP4
HSC
stHSC
6881 cells
UMAP 2
UMAP 1
10X stromal cells
MSC-L1
MPr
SEC
mMPr
MSC-L2
AEC
MSC-S
mMPr (CPr1/OPr1)
MPr (CPr2/OPr2)
Not included
Hematopoietic
c
Femoral sections
PBS
AML
Laminin (sinusoids)
dsRed (AML blasts)
b
10X Stroma
10X HSPC
Total
CXCL
IGF
KIT
ANGPT
ADIPO.
CSF
CCL
CX3C
PDGF
TGFβ
EGF
FGF
RANKL
BMP
PTH
VEGF
WNT
APRIL
TRAIL
Outgoing signals
Incoming signals
MSC-L1
MSC-L2
MSC-S
mMPr
MPr
AEC
SEC
HSC
stHSC
MPP2
MPP3
MPP4
EryP
MkP
GMP
CLP
Relative strength
d
Ifnar2
(Smart-seq)
p=6.4x10-8
Irf7
(Smart-seq)
p=0.0003
Log10 norm. counts
MSC-L
MSC-S
mMPr
MPr
e
tdTomato+
Mx1-Cre:Rosa26-mT/mG
pIC
0.125 mg
i.p.
q48h (3x)
GFP+
GFP+
p<0.0001
% of cells
MSC-L
MSC-S
MPr
Mx1-Cre-
Mx1-Cre+
tdTomato
GFP
0
0
0.01
64.3 ±17.2
21.0 ±10.5
2.3 ±1.1

**Extended Data Figure 9 | MSC-L mediate most interactions with hematopoietic cells. a**, UMAPs showing 10 different HSPCs individually isolated by flow cytometry from young untreated mice (n = 2 biological replicates, one with 5 male mice and the other with 5 female mice, 2 technical replicates per sample) and oligo-hashed before 10X scRNA-seq analyses (left) and the reference 10X stroma map (right) with selected clusters for interactome analysis. **b**, Results of CellChat interactome analysis showing significant families of outgoing (top) and incoming (bottom) signaling molecules between stromal populations and HSPCs. **c**, Representative immunofluorescence images of laminin staining for sinusoidal vessels and dsRed fluorescence from MLL/AF9 AML blasts in CM of femoral sections. **d**, **b,** Expression of indicated interferon genes in Smart-seq mesenchymal populations. Data are violin plots of Log10 normalized (norm.) Smart-seq counts with median; *P. values*, Wilcoxon rank sum test. **e**, Schematic showing tdTomato to GFP conversion upon pIC treatment in *Mx1-Cre:Rosa26-mT/mG* lineage tracing mice (top left), with representative flow cytometry plots (bottom) and quantification (top right) of the proportion of cells converting to GFP in the indicated stromal population. Data are means ± S.D. with points showing values for individual mice; *P. values*, one-way ANOVA with Tukey's *post hoc* test.

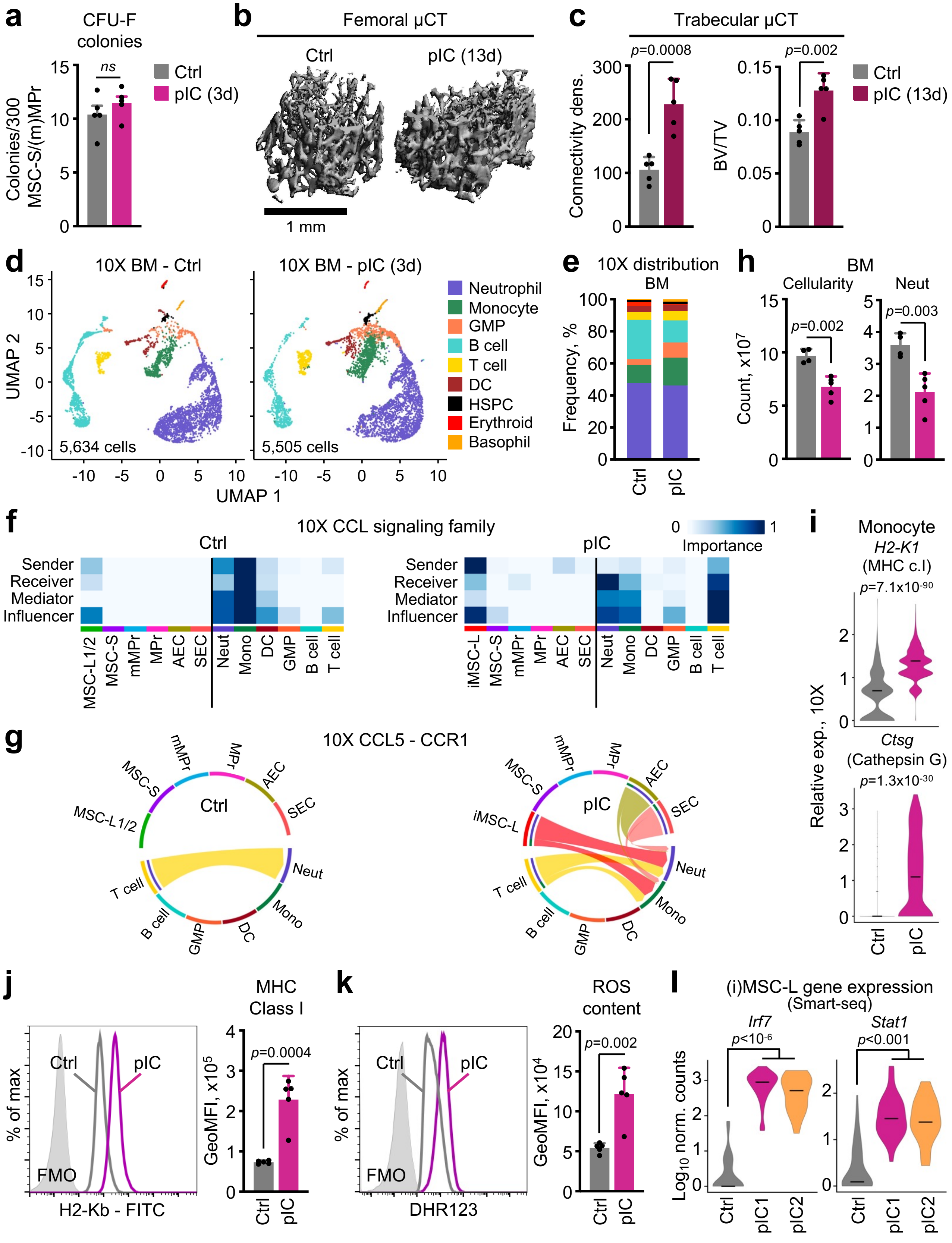
a
CFU-F colonies
Colonies/300 MSC-S/(m)MPr
ns
Ctrl
pIC (3d)
b
Femoral μCT
Ctrl
pIC (13d)
1 mm
c
Trabecular μCT
p=0.0008
p=0.002
Connectivity dens.
BV/TV
Ctrl
pIC (13d)
d
10X BM - Ctrl
10X BM - pIC (3d)
UMAP 2
UMAP 1
5,634 cells
5,505 cells
Neutrophil
Monocyte
GMP
B cell
T cell
DC
HSPC
Erythroid
Basophil
e
10X distribution BM
Frequency, %
Ctrl
pIC
h
BM
Cellularity
Neut
p=0.002
p=0.003
Count, x10^7
f
10X CCL signaling family
Ctrl
pIC
Importance
Sender
Receiver
Mediator
Influencer
MSC-L1/2
MSC-S
mMPr
MPr
AEC
SEC
Neut
Mono
DC
GMP
B cell
T cell
iMSC-L
i
Monocyte
H2-K1
(MHC c.I)
p=7.1x10-90
Ctsg
(Cathepsin G)
p=1.3x10-30
Relative exp., 10X
Ctrl
pIC
g
10X CCL5 - CCR1
Ctrl
pIC
j
MHC Class I
Ctrl
pIC
FMO
% of max
H2-Kb - FITC
p=0.0004
GeoMFI, x10^5
k
ROS content
DHR123
p=0.002
GeoMFI, x10^4
l
(i)MSC-L gene expression
(Smart-seq)
Irf7
p<10-6
Stat1
p<0.001
Log10 norm. counts
pIC1
pIC2

**Extended Data Figure 10 | iMSC-L modulate monocyte dynamics in the BM niche. a**, Quantification of colony forming unit-fibroblast (CFU-F) obtained from 300 MSC-S/(m)MPr isolated from mice injected with PBS (Control, Ctrl) or pIC for 3 days (3d). **b-c,** Representative micro-computed tomography images from central femurs (b) and quantification of trabecular bone connectivity density and bone volume/total volume (BV/TV) (c) in mice injected with PBS (Ctrl) or pIC for 13 days (13d). **d-e,** 10X scRNA-seq profiling of BM cells isolated from control (n = 1, 2 female mice) and 3 day-pIC-injected (n = 1, 1 male & 1 female mouse) mice: (d) UMAP of merged datasets with number of analyzed cells; and (e) frequency of identified BM cell types. DC: dendritic cells. **f-g,** Heatmaps showing major participants in CCL family signaling interactions (f) and Chord plots showing predicted senders and receivers for CCL5-CCR1 interactions (g) between indicated mature BM and stromal cell types (dataset from **Fig. 5g**) identified by CellChat analyses of 10X scRNA-seq datasets. **h,** Cellularity (left) and neutrophil counts (right) in the BM of control and 3 day-pIC-injected mice. **i,** Expression of activation genes in monocytes cluster from 10X BM dataset. Results are shown as violin plots of relative expression (exp.) of 10X SCT transformed counts with median. **j-k,** Representative flow cytometry plots (left) and quantification (right) of MHC class I (H2-Kb) expression (j) and DHR123 expression estimating reactive oxygen species (ROS) content (k) in monocytes of PBS or 3 day-pIC-injected mice, with fluorescence minus one (FMO) control. Results are expressed as geometric mean fluorescence intensity (GeoMFI). **l,** Expression of *Irf7* and *Stat1* genes in indicated Smart-seq (i)MSC-Ls. Results are shown as violin plots of Log10 normalized (norm.) Smart-seq counts with median. Data in (a), (c), (h), (j), and (k) are means ± S.D. with points showing values for individual mice; *P. values*, Student's t test. *P. values* in (i) and (l) are Wilcoxon rank sum test.

**Extended Data Table 1 | Distribution of stromal and hematopoietic cells in bone marrow spatial compartments.** Distribution of cells in Louvain clusters in 10X stromal map shown in Fig. 1c, with origin of cells in either central marrow or endosteal samples. Top 50 most significantly upregulated genes in each cluster compared to all other clusters.

*(separate Excel file)*

**Extended Data Table 2 | Key gene markers in Smart-seq populations.** Top 100 most significantly upregulated genes in each stromal population compared to all other cell types.

*(separate Excel file)*

**Extended Data Table 3 | Gene signatures used for annotation of MSCs in 10X mesenchymal atlas**

| Cell | $LepR^+$ MSC | Sca-$1^+$ MSC |
|---|---|---|
| Genes | *Lepr* | *Ly6a* |
| | *Ibsp* | *Cd34* |
| | *Mgp* | *Thy1* |
| | *Ogn* | *Mfap5* |
| | *Cxcl12* | *Gsn* |
| | | *Clec3b* |

Genes used for annotation of mesenchymal stromal cell (MSC) populations in 10X mesenchymal atlas.
