## Supplemental Figures 1 to 3 for "Inflammation perturbs hematopoiesis by remodeling specific compartments of the bone marrow niche"

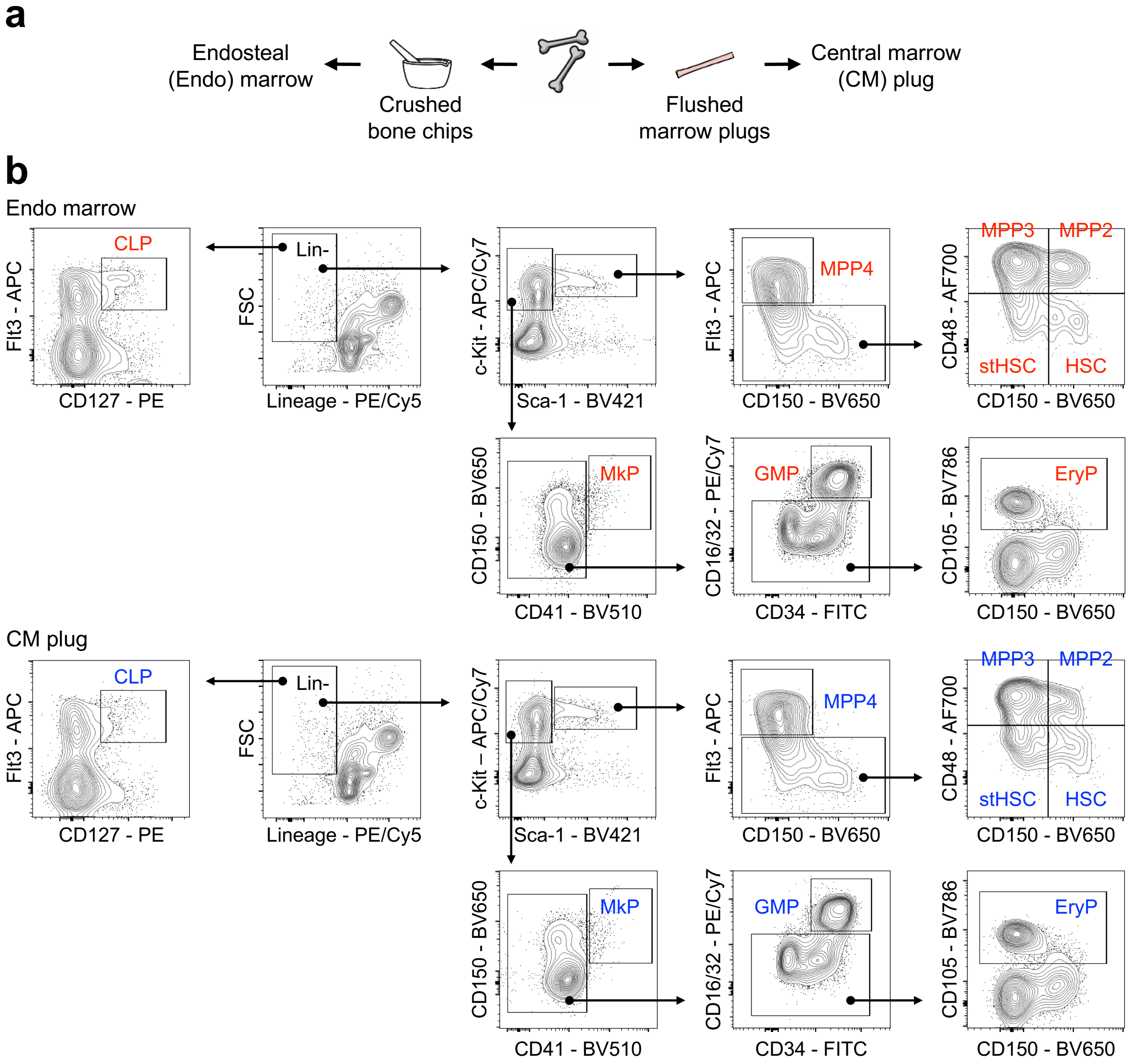


**Supplementary Information 1 | BM HSPC gating strategy. a**, Scheme showing the isolation of CM plug from flushed bones and Endo marrow from crushed bones. **b**, gating scheme for the indicated HSPC populations in Endo marrow (top) and CM plug (bottom). HSC: hematopoietic stem cell; stHSC: short-term HSC; MPP2/3/4: multipotent progenitor 1/2/3; MkP: megakaryocyte progenitor; CLP: common lymphoid progenitor; GMP: granulocyte macrophage progenitor; EryP: erythroid progenitor.

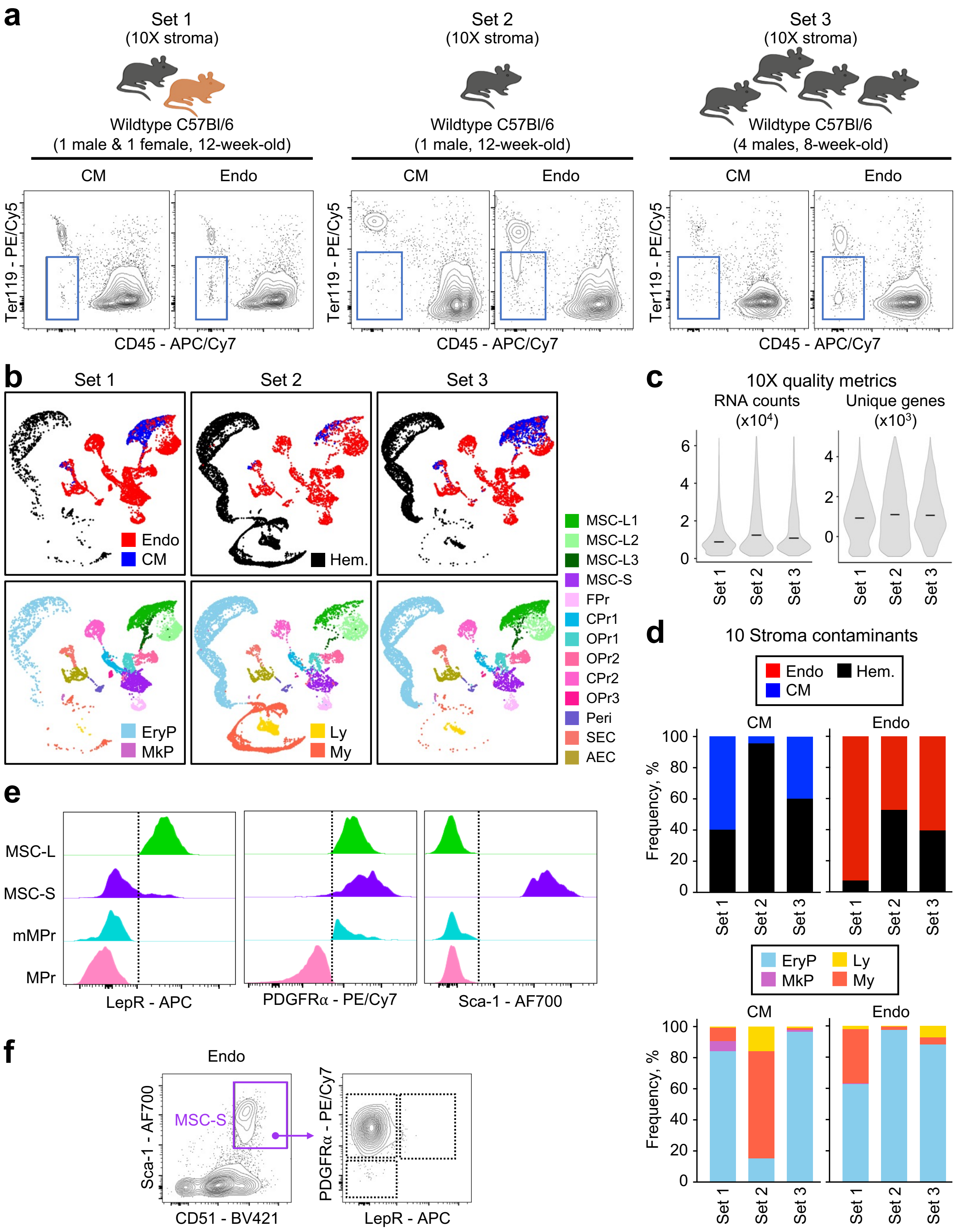

a
Set 1
(10X stroma)
Wildtype C57Bl/6
(1 male & 1 female, 12-week-old)
Set 2
(10X stroma)
Wildtype C57Bl/6
(1 male, 12-week-old)
Set 3
(10X stroma)
Wildtype C57Bl/6
(4 males, 8-week-old)
CM
Endo
Ter119 - PE/Cy5
CD45 - APC/Cy7
b
Set 1
Set 2
Set 3
Endo
CM
Hem.
EryP
MkP
Ly
My
MSC-L1
MSC-L2
MSC-L3
MSC-S
FPr
CPr1
OPr1
OPr2
CPr2
OPr3
Peri
SEC
AEC
c
10X quality metrics
RNA counts (x10^4)
Unique genes (x10^3)
d
10 Stroma contaminants
Endo
Hem.
CM
Frequency, %
EryP
Ly
MkP
My
e
MSC-L
MSC-S
mMPr
MPr
LepR - APC
PDGFRα - PE/Cy7
Sca-1 - AF700
f
Endo
MSC-S
Sca-1 - AF700
CD51 - BV421
PDGFRα - PE/Cy7
LepR - APC

**Supplementary Information 2 | Generation of the integrated reference 10X stroma map and additional stromal cell flow characterization. a**, Scheme showing the 3 independent biological replicate sets used for flow-isolation of CM and Endo Ter119$^-$/CD45$^-$ stromal cells for 10X scRNA-seq analyses. **b,** UMAP dimension reductions showing for each replicate set the CM (blue) or Endo (red) origin of the cells, with hematopoietic cell (Hem.) contaminants (top) and naming of stromal cell clusters (bottom). AEC: arterial endothelial cells, SEC: sinusoidal endothelial cell, MSC-S: Sca-1$^+$ mesenchymal stromal cell (MSC), OPr: osteoblast progenitor, CPr: chondroblast progenitor, FPr: fibroblast progenitor, MSC-L: leptin receptor (LepR)$^+$ MSC; EryP: erythroid progenitor, MkP: megakaryocyte progenitor, Ly: lymphoid cells; My: myeloid cells (together with rare HSPC). **c,** Violin plots showing RNA counts and number of unique genes observed in each replicate set, with medians. **d,** Bar plots showing the amount (top) and type (bottom) of contaminating hematopoietic cells in each replicate set. **e,** Representative flow cytometry plots showing expression of the indicated markers in indicated stromal cell populations. (m)MPr: (multipotent) mesenchymal progenitor. **f**, Representative flow cytometry plots showing lack of LepR expression in Endo MSC-S.

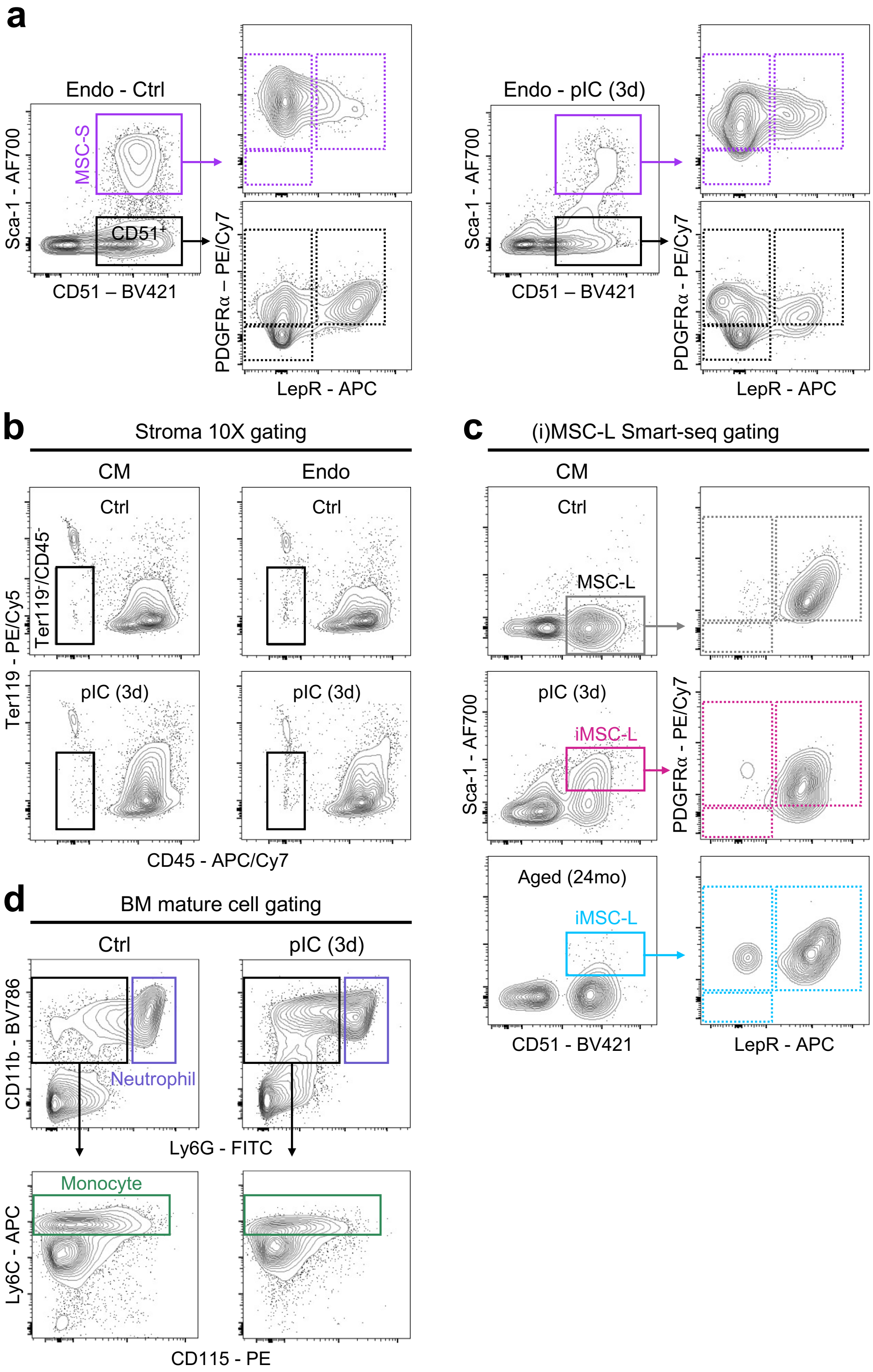
a
Endo - Ctrl
MSC-S
CD51+
Sca-1 - AF700
CD51 – BV421
PDGFRα – PE/Cy7
LepR - APC
Endo - pIC (3d)
Sca-1 - AF700
CD51 – BV421
PDGFRα - PE/Cy7
LepR - APC
b
Stroma 10X gating
CM
Endo
Ctrl
Ctrl
pIC (3d)
pIC (3d)
Ter119 - PE/Cy5
Ter119-/CD45-
CD45 - APC/Cy7
c
(i)MSC-L Smart-seq gating
CM
Ctrl
MSC-L
pIC (3d)
iMSC-L
Aged (24mo)
iMSC-L
Sca-1 - AF700
CD51 - BV421
PDGFRα - PE/Cy7
LepR - APC
d
BM mature cell gating
Ctrl
pIC (3d)
CD11b - BV786
Neutrophil
Ly6G - FITC
Monocyte
Ly6C - APC
CD115 - PE

**Supplementary Information 3 | Detailed flow cytometry analyses and gating schemes. a**, Representative flow cytometry plots showing unchanged marker expression in Endo MSC-S and $CD51^{+}$ stroma cells in 3 day-pIC-injected mice. **b-c,** Flow cytometry gating schemes used for isolation of CM and Endo $Ter119^{-}/CD45^{-}$ stromal cells from 3 days (3d) PBS control (Ctrl) or pIC-treated mice for 10X scRNA-seq analyses (b), and single (i)MSC-L cells for Smart-seq analyses (c). Aged (24mo): 24-month-old mice. **d,** Flow cytometry gating scheme used for identification of mature myeloid cells in the BM.
